# A Joint Tau Propagation and Neuroinflammation Model Reinforces Inflammatory Modulation of Network-Driven Spread in Alzheimer’s Disease

**DOI:** 10.64898/2026.01.07.698214

**Authors:** Ishaan Dhaliwal, Robin Sandell, Ashish Raj

## Abstract

Alzheimer’s disease (AD) is a progressive neurodegenerative disorder and the leading cause of dementia. Despite over a century of research, AD remains untreatable due to an incomplete understanding of its underlying mechanisms. While amyloid beta has dominated therapeutic efforts, tau pathology and neuroinflammation represent critical disease drivers and intriguing therapeutic targets. We developed an extended computational model building upon the open-source Aggregation Network Diffusion (AND) framework by coupling spatial tau propagation and aggregation with ordinary differential equations representing key inflammatory cascades. The model incorporates M1/M2 microglia and astrocytic activation, cytokine-mediated feedback loops, and neuronal loss, all modulated via region-specific genetic expression matrices of ApoE and TREM2 with variant-specific weighting. The model maintains biologically plausible tau dynamics while generating robust inflammatory marker trends, serving as a computational testbed for hypothesis generation and mechanistic exploration. The inflammatory components independently capture experimental observations and display a marked M1/M2 microglia divergence. The model demonstrates enhanced similarity to regional tau propagation patterns due to inflammatory and genetic components, reinforcing neuroinflammation’s role in tau spread and highlighting the need to incorporate these processes in modeling efforts. Finally, through parsimony analysis, we identify microglia and pro-inflammatory rates (including microglia-facilitated tau spread) as key contributors to improved model accuracy, informing future modeling approaches.

## Introduction

Alzheimer’s disease (AD) is the most common neurodegenerative disorder (1) affecting approximately 7 million Americans—a number expected to reach 13 million by 2060 (Alzheimer’s Association, 2024). First reported by Alois Alzheimer (1906), AD pathology is characterized by amyloid beta (Aβ) aggregation in neuritic plaques, tau protein accumulation in neurofibrillary tangles, and sustained neuroinflammation through chronic immune pathway activation (3, 4, 5). These processes impair neuronal function, triggering progressive cognitive decline and memory loss. Despite clear clinical symptoms, the specific mechanisms and temporal sequence of disease mediators remain poorly understood, significantly undermining treatment development (6).

Tau, a microtubule-associated protein (MAP) abundant in neurons, stabilizes and regulates microtubules by binding tubulin dimers (7). In pathological conditions, hyperphosphorylation disrupts tau’s normal function and promotes aggregation (8).

Misfolded tau forms soluble oligomers that develop into neurofibrillary tangles, a hallmark of neurodegenerative diseases including AD. Tau pathology spreads through a “prion-like” process: after secretion, misfolded tau is taken up by functionally or anatomically connected neurons via endocytosis and receptor-mediated pathways (9, 10, 11), seeding native tau misfolding and propagating neurodegeneration.

Given the complex and network-centric interplay between key pathologies, researchers have increasingly turned to computational models to address gaps arising from purely experimental in vivo mouse or human systems. For tau specifically, microscopic aggregation models include the nucleated polymerization model (12), heteromer model (13), and modified Smoluchowski coagulation equations (14). Macroscopically, tau propagation modeling employs connectome-based networks (15), epidemic spreading-based propagation (16), and graph theory-derived diffusion (17). These works have culminated in more detailed models, including advancing network-mediated spread to co-evolving Aβ and tau species (18), incorporating axonal transport into directional spread (19), and using modified graph neural networks (20).

Parallel to tau aggregation and spread, AD features sustained inflammatory responses. Pathological tau and Aβ trigger neuroinflammation by activating glial cells, including microglia and astrocytes (21). In a feedback loop, glial activation produces signaling molecules, notably cytokines—small proteins that stimulate or inhibit microglial activation and inflammation (22). Continuous presence of aggregated proteins shifts balanced inflammatory responses toward chronically activated microglia. Pro-inflammatory cytokines amplify activation and proliferation of other microglia through factors like GM-CSF and M-CSF (23) while exacerbating tau pathology by inducing neuronal stress. Cytokines activate intracellular signaling pathways including glycogen synthase kinase-3 beta (GSK-3β) (24) and p38 mitogen-associated protein kinase (MAPK), driving tau hyperphosphorylation. This creates a self-perpetuating loop: misfolded tau activates microglia, leading to cytokine release that amplifies tau aggregation and facilitates spread, further activating microglia. Over time, this cycle drives chronic neuroinflammation, neuronal damage, and synaptic dysfunction, accelerating neurodegeneration.

Pathologic and inflammatory processes are mediated by key genetic modulators including TREM2 and APOE, with variants serving as prominent AD risk factors. TREM2 encodes a microglial receptor crucial for inflammatory responses with complex roles in AD progression. While TREM2 does not directly bind tau, it plays important roles in tau clearance early in disease progression (25) and its IL-4 production limits excessive inflammation (26 However, particularly later in AD, TREM2 triggers activation of extracellular signal-regulated kinase (ERK) (27) and phosphoinositide 3-kinase (PI3K) (28) pathways, promoting pro-inflammatory cytokines and tau misfolding. TREM2 deficiency or loss-of-function mutations significantly promote tau aggregation and disease pathology, with studies showing the R47H mutation increases tau propagation and pro-inflammatory cytokines while decreasing misfolded protein clearance (29). APOE is similarly involved in immune responses and mediates lipid transport in the brain. The APOE4 variant is the strongest genetic risk factor for AD (30), displaying significantly impaired clearance abilities that exacerbate amyloid plaque accumulation. APOE4 increases tau phosphorylation in murine models (31).

However, the use of computational modeling for incorporating key inflammatory factors and mediators remain in its infancy. This is a significant gap, because modeling would appear to be perfectly suited to interrogate the multi-factorial, complex and mutually interacting processes described above. Such processes cannot be fully studied using purely experimental in vitro or in vivo models, such as transgenic animals and human neuroimaging. Animal models, while valuable, often fail to replicate human tau pathology complexity and may not fully capture clinical variability, sometimes producing inaccurate results. Similarly, neuroimaging techniques provide important information about tau deposition and brain atrophy patterns but cannot directly link tau propagation to molecular mechanisms (32). While several computational models exist for neuroinflammation in AD, including those by Hao and Friedman (2016) and Puri and Li (2010), these typically apply at specific brain locations or globally (33, 34). Such approaches capture temporal evolution of inflammation and protein pathology but generally do not account for spatial heterogeneity across brain regions, nor the key role of network spread.

Despite tau and neuroinflammation’s roles, AD research has historically focused on amyloid pathology, driven by the amyloid cascade hypothesis which posits that upstream Aβ accumulation triggers tau pathology and downstream neurodegeneration. However, most Aβ-targeting drugs have failed to significantly alter AD pathology (35). Disconnects between the temporal and regional spread of Aβ and tau suggest other mechanisms are involved. Studies show stronger spatial relationships between tau and AD progression (36), with tau pathology also prominent across other neurodegenerative diseases (37).

Thus, this study presents a computational model integrating tau aggregation and network-based propagation with AD-specific neuroinflammation dynamics, emphasizing inflammatory pathways and intermediates. Our goal is to fill the gap between current computational modeling of tau pathology and the complex manner in which inflammatory processes mediate those processes. In contrast to prior models of inflammation, our approach encompasses all brain regions and network connections simultaneously and allows for regionally varying patterns of both mediating factors as well as the resulting tau pathology distributions. While Aβ is an important modulator, we omit it to focus on non-Aβ-informed therapies. The proposed model builds upon a modified AND (Aggregation Network Diffusion) framework (38) to simulate tau dynamics tuned to reflect AD dynamics. We chose AND as our base model because it already incorporates a broad set of processes—local production and aggregation kinetics of tau, from monomers to oligomers to tangles, and network-level diffusion of oligomeric tau to distant regions via the connectome. Hence, we build on the AND model and introduce a wide range of global as well as regionally specific inflammatory systems and their induced dynamics within brain regions. These aspects are challenging to achieve in simpler diffusion-only models like the network diffusion model (NDM; 17) or other network-based spread approaches. Using AND’s simultaneous local and macroscopic approach, we couple it with ordinary differential equations (ODEs) describing interactions among tau, microglia, cytokines, astrocytes, and additional AD-relevant elements within brain regions. Relationships and parameterizations for these interactions are informed by experimentally established mechanisms and utilize existing computational relations, including the PDE model by Hao and Friedman (33).

The model, which we call the Aggregation Network Diffusion with Inflammation (ANDI) model, was assessed on empirical tau PET data from a study at Yonsei University (39). In the end, we found not only that the inflammation-enhanced model improved on the base model’s empirical fits, but also that it was able to remove spatial inconsistencies observed in previous network-based models. We also found that not all inflammatory actors exert a strong role in governing tau spread; some, like microglia are important, while others appear less so. By incorporating spatially variable inflammatory systems into network models of tau, the proposed model represents a step change in the modeling community, providing novel insights that can inform future experimentalists and may have broad applicability and translational potential in other related disorders.

## Methods

### Base Model

To construct the initial tau aggregation and propagation model, we used the data and framework from the Raj et al. (2021) AND model. The model creates a network with vertices representing brain regions and simulates aggregation within each node. First, the model establishes generation of misfolded tau monomers from healthy tau, assuming spontaneous production in a gamma distribution pattern, governed by the equation below where represents the ith vertex or node of the imposed connectome, is the seeding region vertex, t is time, and α is a constant.

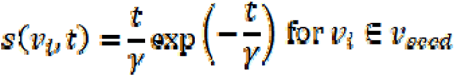

Using the monomer concentration as a source term, the model then simulates aggregation from monomer to oligomer to fibril using a modified discrete Smoluchowski coagulation equation (capped at M, an arbitrary point where aggregates are considered tangles). For the initial seeding region, the equation is defined as:

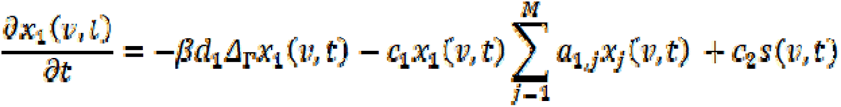

The first term represents diffusion (described later), the second term represents aggregation into larger complexes, and the third term represents monomer generation. Here, represents the concentration of tau of length m at vertex v at time t. α and β are aggregation and monomer production coefficients, and all “a” constants are rate constants. For other regions, oligomer aggregation at lengths less than M is governed as follows:

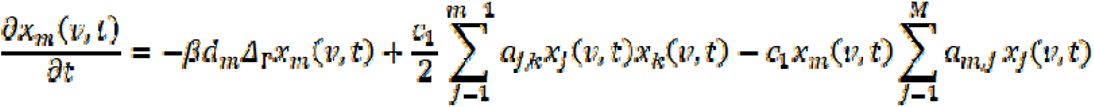

where j and k are two polymers aggregating to form m. In both Equations 2 and 3, the first term represents diffusion, explained later. Finally, tangle aggregation is described by a distinct equation without terms for diffusion or fragmentation.

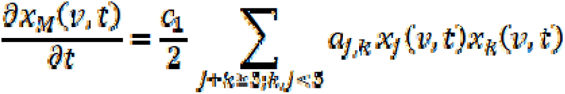

The next step integrates propagation features. As outlined in the AND model, a reaction diffusion model combined with graph theory attributes is utilized. The model relies on a hypothetical brain network with nodes representing compartmentalized gray matter structures (from MRI). The network’s edges represent white matter fiber pathways determined using connection strengths. We model tau diffusion in this imposed network. For example, given two nodes, the change in tau concentration at one node equals diffusion to that node minus diffusion away from it. In a larger network, the rate of change of tau at one node is proportional to the total connectivity and tau concentration at each node. This can be simplified as the Laplacian matrix of the vector describing tau concentration at each node. Adding rate constant D and diffusivity constant for the mth oligomer , we reach the diffusion term used in Equations 2 and 3 to describe aggregates at each node.

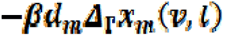

### Adding Inflammation Module

To add neuroinflammatory capabilities, we built upon the existing AND model by coupling it with ordinary differential equations (ODEs). This system encompassed state variables for monomeric tau, intracellular tau tangles, extracellular tau tangles, healthy neuron population, dead neuron population, M1 microglia, M2 microglia, pro-inflammatory cytokines (collectively represented by TNF-alpha), anti-inflammatory cytokines (collectively represented by IL-10), astrocytes, and TGF-beta (Figure 1). The inflammatory equations were adapted from the model developed by Hao and Friedman (2016), modified to ODEs rather than partial differential equations, simplified to fewer state variables, and changed so the existing tau dynamics were replaced by the AND model.

**FIG 1:**
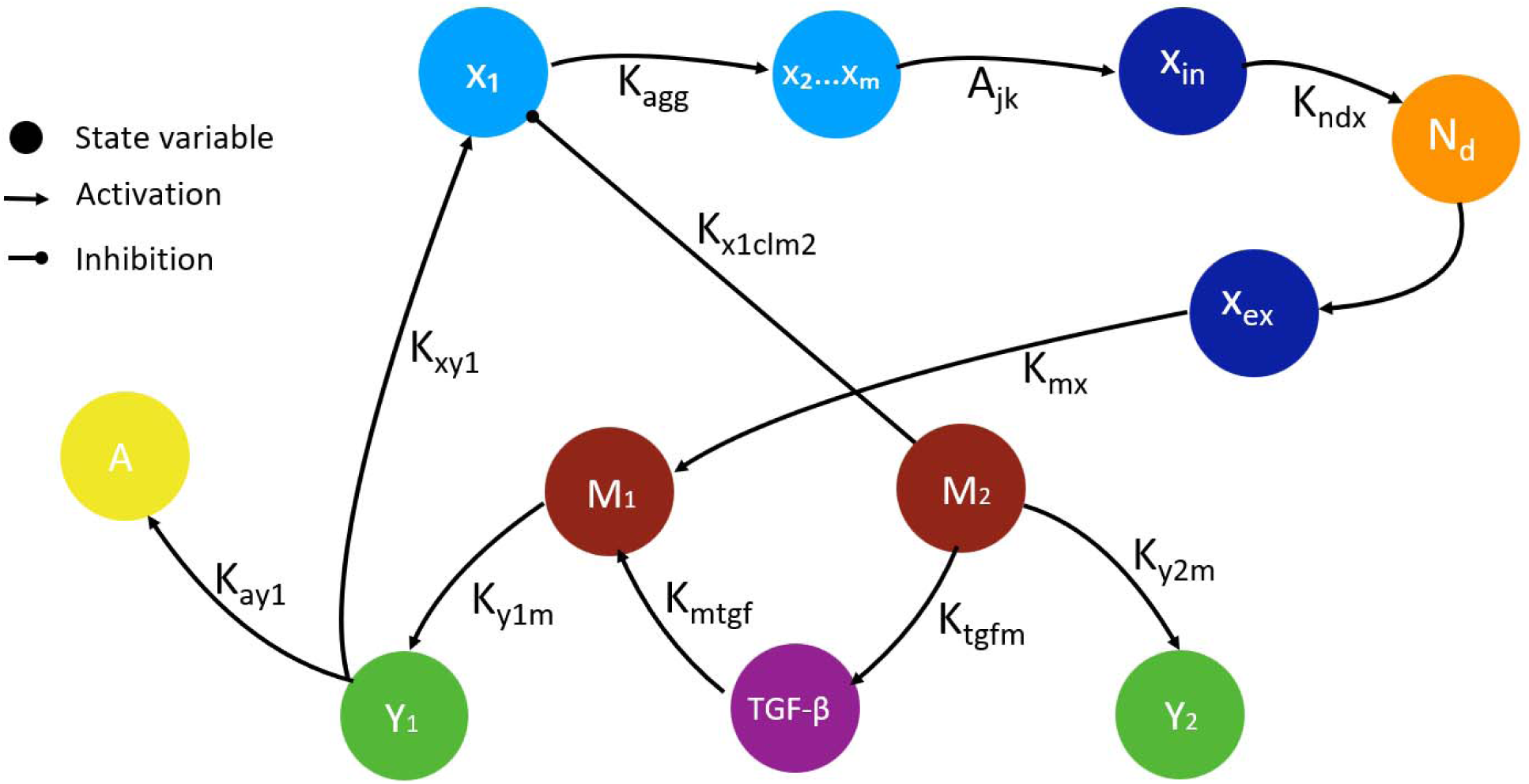
Visualization of the key interactions in the inflammatory portion of the model. Circles represent state variables, and arrows indicate parameters. Certain parameters (including clearance rates and parameters that impact tau propagation) are not included for the sake of simplicity.

Specifically, the model uses tau monomer and tangle levels from the AND model to simulate extracellular neurofibrillary tangles, which are released upon neuron death, and are cleared at a rate dependent on concentration.

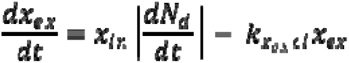

Intracellular NFTs cause neuronal death through microtubule destabilization, modeled using a Michaelis-Menten relationship. Cytokines influence neuronal death similarly, with pro-inflammatory cytokines contributing to death and anti-inflammatory cytokines protecting neurons.

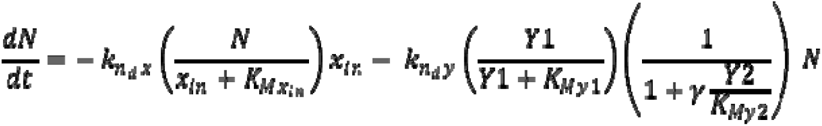

The differential equation for dead neurons is the opposite of the healthy neuron equation with an additional clearance term dependent on microglia populations.

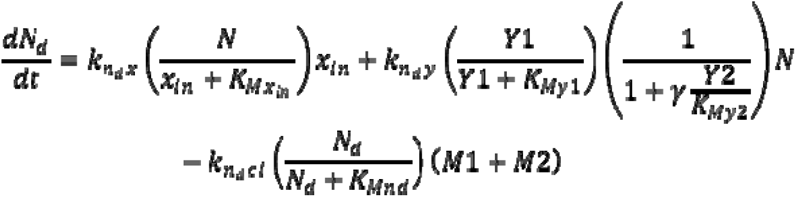

Astrocytes are activated primarily by pro-inflammatory cytokines (TNF-alpha), providing the following equation.

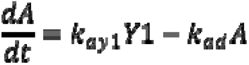

Microglia were modeled as either M1 (pro-inflammatory) or M2 (anti-inflammatory). The transition rate between phenotypes is governed by TGF-beta, which modulates microglia from M1 to M2 phenotype. Both types are activated by extracellular NFTs and their respective cytokines. For cytokine-based activation, the model utilizes an epsilon term governing non-linear cytokine growth, as in Hao and Friedman (2016). M1 microglia experience cytokine activation from the ratio of epsilon1 (based on pro-inflammatory cytokines) to total cytokine concentration multiplied by beta, a parameter reflecting TNF-alpha’s higher potency compared to IL-10. Similarly, M2 microglia are activated by cytokines via the term epsilon2/(repsilon1+epsilon2), reflecting the ratio of anti-inflammatory to total cytokine concentration.

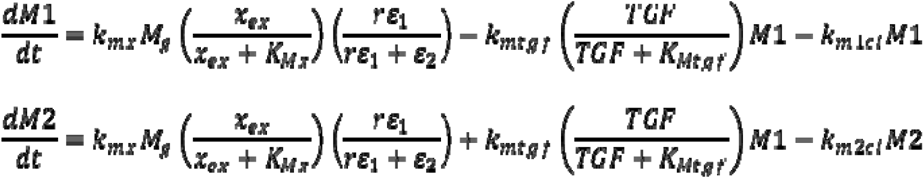

TGF-beta modulates M1 to M2 microglia, and exhibits a positive feedback loop with M2 microglia, modeled below.

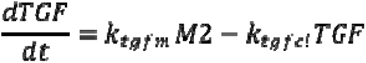

Pro-inflammatory cytokines (referred to as C1) and anti-inflammatory cytokines (referred to as C2) are produced by M1 and M2 microglia respectively and cleared depending on concentration.

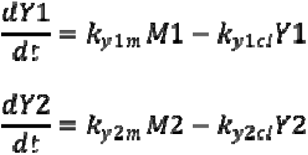

To integrate with the tau portion of the model, an additional term was added to the monomeric tau differential equation based on pro-inflammatory cytokine levels. This term uses a Hill equation with a Hill coefficient of 2 simulating positive cooperativity between pro-inflammatory cytokines and tau misfolding, yielding the following equation.

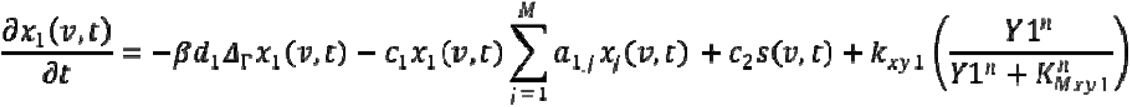

Finally, feedback from the tau portion substitutes into intracellular NFT levels in the inflammation portion, completing the bidirectional loop. The state variables and parameters mentioned above are summarized in Table 1 and Table 2.

**Table 1:** List of time-dependent state variables in the model and their interpretation.

| State Variable | Symbol |
| --- | --- |
| Astrocytes | A |
| Oligomeric tau (size m) | $x_m$ |
| Intracellular neurofibrillary tangles | $x_{in}$ |
| Extracellular neurofibrillary tangles | $x_{ex}$ |
| Healthy neurons | N |
| Dead neurons | $N_d$ |
| Microglia (M1 or M2) | M1, M2 |
| TGF-beta | TGF |
| Cytokines (pro-inflammatory or anti-inflammatory) | Y1, Y2 |

**Table 2:**
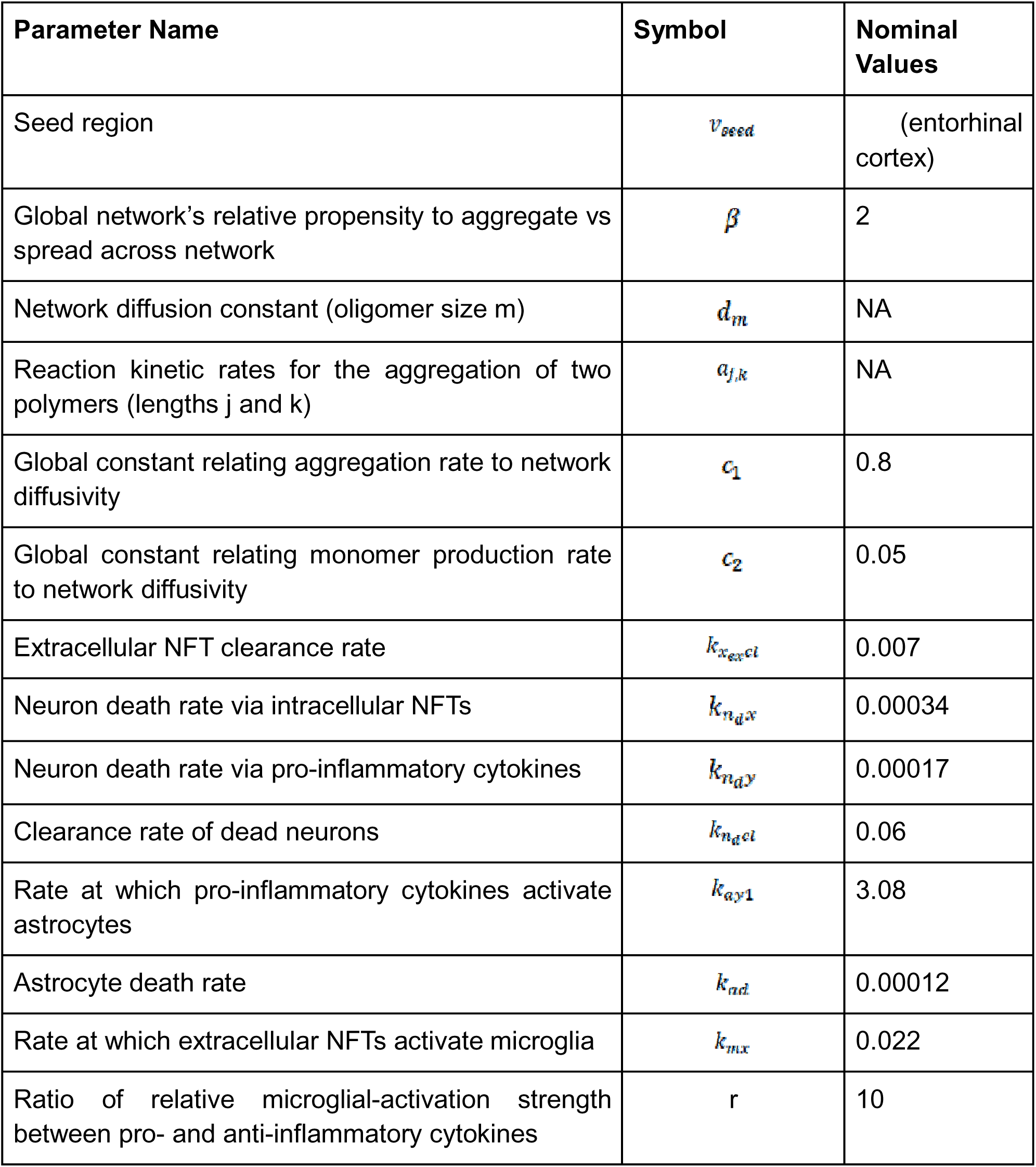
Glossary of model parameters, their biological interpretation, and their nominal values.

### Genetic Isoform-specific Modeling

To simulate TREM2 and APOE genetic pathway influence, we incorporated brain region-specific gene expression data into the model. This data constructed an expression matrix capturing relative abundance and spatial distribution of these genes across neural environments. The matrix was then applied to modulate key parameter rate constants within the differential equation system, emulating regulatory impacts these genetic factors exert on pathological processes including neuroinflammation, neuronal survival, and protein aggregation.

To capture genetic heterogeneity, each expression matrix was scaled by isoform-specific weight constants. For APOE, this included allelic variants ApoE2, ApoE3, and ApoE4, which differentially influence neurodegenerative disease risk. Similarly, for TREM2, separate weightings were introduced for wild-type TREM2 and the disease-associated R47H mutation. By incorporating these weighted matrices, we captured both regional gene expression differences and genotype-dependent modulation.

### Data sources and processing

Tau-PET data used in this study was obtained from the public previously published study at Yonsei University, South Korea, detailed in a separate paper (39). Empirical 18F-AV1451 imaging data was obtained from 128 consecutive patients consisting of 53 patients with probable Alzheimer’s dementia, 52 patients with amnestic mild cognitive impairment (aMCI), and 23 patients with nonamnestic MCI (naMCI) as well as 67 matched controls. Specifically, subjects were intravenously injected with 281.2MBq of Flortaucipir for tau PET, and, at 80 minutes after the injection, PET images were taken. From here, PET images underwent an ordered subsets expectation maximization algorithm in a 256 x 256 x 223 matrix with 1.59 x 1.59 x 1 mm voxel size. Additionally, axial T1-MRI was obtained with 3D spoiled gradient with specific parameters: repetition time of 8.28 ms, echo time of 1.6 to 11.0 ms, 20 degree flip angle, 512 x 512 matrix size, 0.43 x 0.43 x 1 mm voxel size in a 3T MR scanner (Discovery MR750; GE Medical Systems).

To establish regional PET uptake, the following method was used. First, to establish a reference point, PET images from healthy controls were utilized to generate an average image, which was then normalized to the MNI space using SPM8’s linear and non-linear transformations. This transformation was then consistently applied to all individual PET images, aligning them with the same voxel resolution as the 86-region DK atlas. Finally, mean AV1451 values were computed for each of the GM ROIs to quantify tau-PET signals within specific brain regions.

### Diffusion MRI (dMRI) and tractography processing

All processing was carried out within a custom pipeline based on the NiPype framework (40). T1 images were segmented into gray (GM), white matter (WM), and CSF tissue maps using SPM, where T1 images were registered and transformed to MNI space. dMRI volumes were corrected for Eddy currents and small head movements by registering the diffusion-weighted volumes to the first non-diffusion-weighted volume via an affine trans\-formation using FSL FLIRT (41). Skull-stripping was performed using FSL BET. Details regarding this processing can be found in a previous publication (42).

### Structural connectivity network

Different structural connectivity networks were constructed using the same DK parcellations as above. First, we obtained publicly available dMRI data from the MGH-USC Human Connectome Project (HCP) to create an average template connectome. The HCP contains data from 418 healthy brains (43). Subject-specific structural connectivity was computed on dMRI data as described in Abdelnour et al. (44) and Owen et al. (45). *Bedpostx* was used to determine the orientation of brain fibers in conjunction with FSL FLIRT (41). Tractography was performed using *probtrackx2* to determine the elements of the adjacency matrix. 4,000 streamlines were initiated from each seed voxel corresponding to a cortical or subcortical GM region and the number of streamlines reaching a target GM region was recorded. The weighted connection between the two structures, c(i, j), was defined as the number of streamlines initiated by voxels in region i that reached any voxel in region j, normalized by the sum of the source and target region volumes. The average connection strengths between both directions, (c(i, j) and c(j, i)), formed the undirected edges. To determine the geographic location of an edge, the top 95% nonzero voxels were computed for both edge directions and the consensus edge was defined as the union between both post-threshold sets. Details regarding this processing can be found in Verma et al (46).

### Regional Gene Expression Data

Brain region-specific gene expression data for APOE and TREM2 were obtained from the Allen Human Brain Atlas (http://human.brain-map.org/). Expression microarray data were extracted and normalized for each of the 86 Desikan-Killiany (DK) atlas regions, providing quantitative measures of regional gene abundance. The data matrix contained normalized expression values (0-1 scale) for 100 AD-related genes, including APOE and TREM2, across all cortical and subcortical regions. To account for genetic heterogeneity, region-specific expression values for APOE and TREM2 were scaled by isoform-specific weight constants. For APOE, separate scaling factors were applied for E2, E3, and E4 allelic variants based on their differential association with AD risk (30). For TREM2, distinct weightings were used for wild-type TREM2 and the R47H loss-of-function mutation (47). These weighted expression matrices were incorporated into the model to modulate inflammatory and tau-related rate parameters within each brain region, enabling region- and genotype-specific simulation of disease dynamics.

## Results

### Parameter Fitting and Optimization

To tune model parameters, we ran an optimization routine. Using empirical data from regional tau PET scans, we computed the R-value and recorded the peak correlation. The univariate grid search changed key model parameters (Table 2) one at a time while keeping others constant. This was performed using a logarithmic scale to capture coarse tuning, then with finer variations ranging from -60% to +60% of the parameter’s value (Figure 2). Though the model is original, most parameters required little tuning, as many were previously validated in the models it was based on. Nevertheless, the univariate grid search results provided a principled way to select final parameter values and ensure optimal model fit.

**FIG 2:**
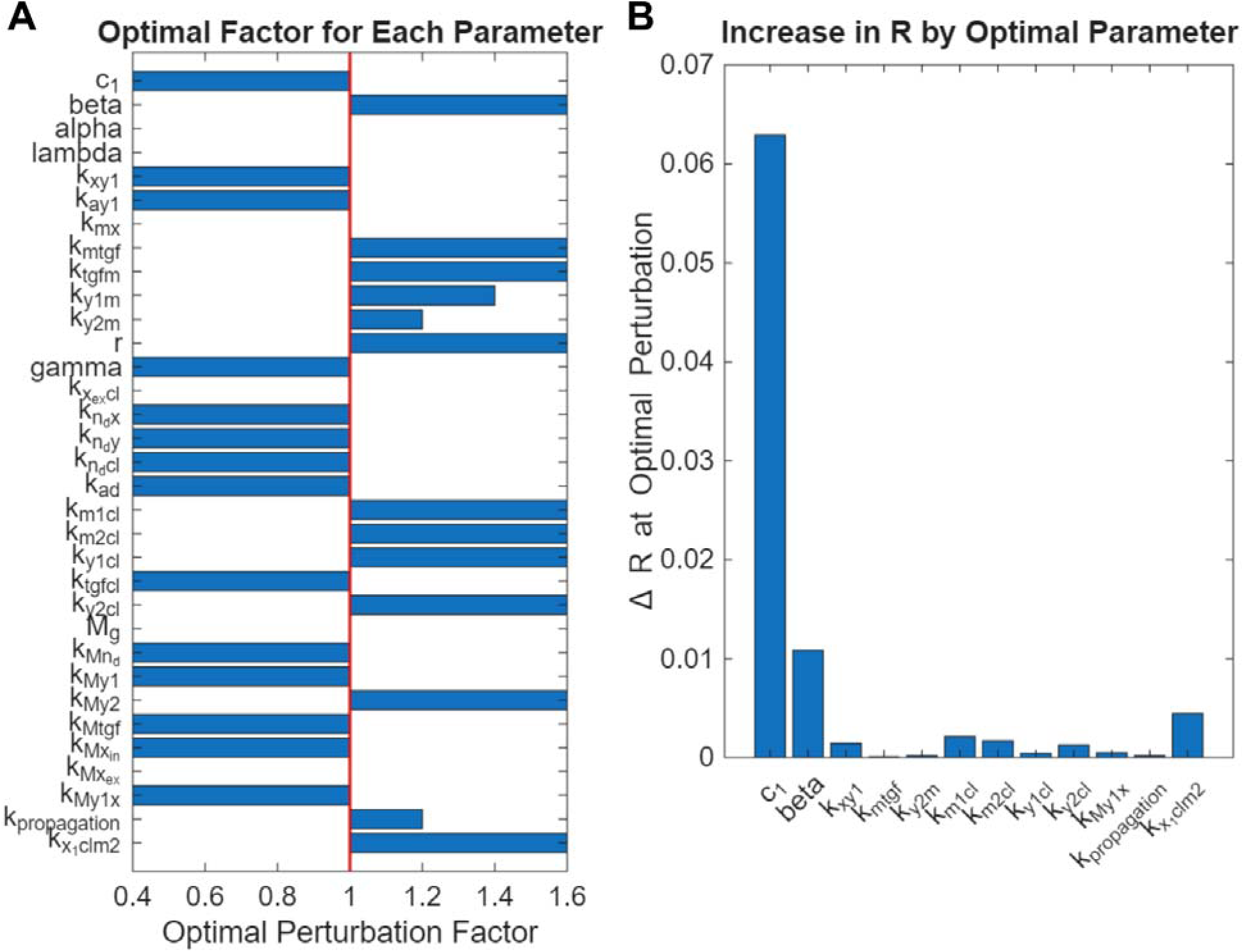
(A) Visualization of the results of the univariate optimization routine, with the displayed analysis tuning parameters within a range of 0.4 and 1.6 of the parameter’s original value. **(B)** Of the optimal changes, the parameters which conferred noticeable differences in R were selected, and their contribution to model fit is visualized.

**FIG 3:**
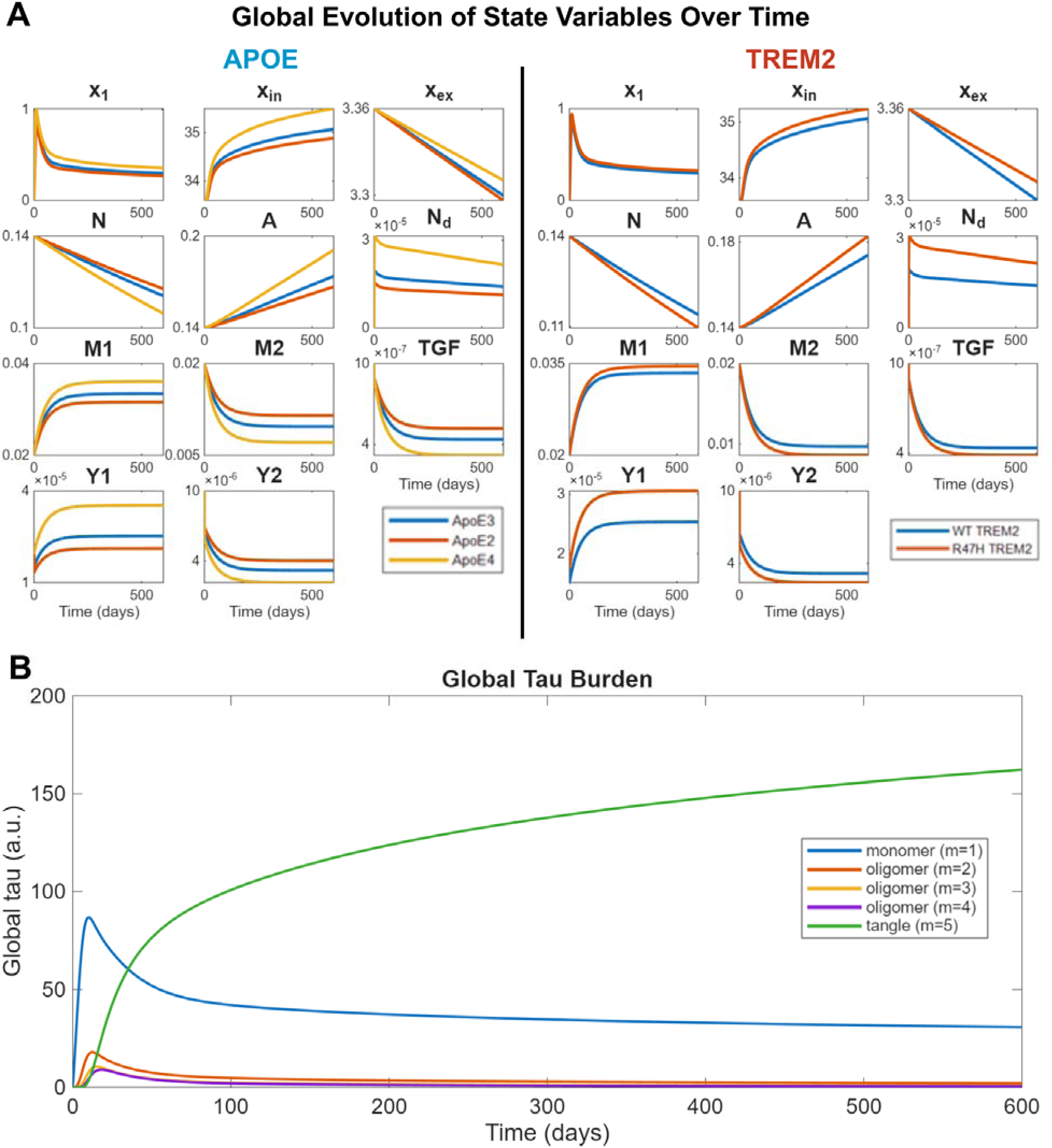
(A) Evolution of the model state variables totaled throughout the brain over time (600 days). The state variables are modulated by region-specific genetic expression, with the impact of ApoE isoforms on the left and TREM2 (wild-type vs R47H) on the right. **(B)** Evolution of tau oligomers (again over the entire brain) using the base model (ApoE3 and wild-type TREM2). Like the AND predecessor, tau length is quantized for convenience and computational load into 5 bins ranging from monomers (m=1) to tangles (m=5) with varying, arbitrary, lengths of oligomers in between.

Of the parameters listed in Panel A and Table 2, only 12 parameters were selected for perturbation (Panel B). Remaining parameters displayed minimal R changes (less than 0.0001) and were omitted. Among perturbed parameters, c_l_ and /3 , both parameters related to diffusion, displayed the most prominent R increase (Panel B). Regarding inflammation- specific parameters, k_xλclm2_ (the tau clearance rate by M2 microglia) most improved model fit. During the earlier logarithmic grid search, the largest parameters adjustments occurred with the aforementioned k_xλclm2_, as well as k_propagation_ (rate of microglia-induced tau spread) and k_xyl_ (rate of tau misfolding mediated by pro-inflammatory cytokines)

### Temporal Evolution of State Variables

Figure 2 shows temporal evolution of the 11 state variables, with modulation via genetic effects. Focusing on inflammation-related state variables, results show characteristic patterns of microglia activation and cytokine dynamics. Specifically, the model demonstrates M1 microglia and pro-inflammatory cytokines increasing with disease progression, with the opposite for M2 microglia and C2 cytokines. Similarly, other state variables display consistent trends, with global neurons decreasing steadily, astrocytes increasing, and TGF-beta decreasing.

Panel A also shows the impact of APOE isoforms and wild type versus mutated TREM2 on the progression of state variables. The E2 isoform is the most neuroprotective, shown by lower monomeric tau and tangles plus elevated M2 microglia and anti-inflammatory cytokines. The E4 isoform increases disease progression with elevated inflammatory responses and suppressed anti-inflammatory factors. Similarly, R47H mutated TREM2 increases neuron death and overall pathology while lowering M2 and anti-inflammatory cytokine responses. Examination of tau dynamics by species size (Panel B) shows that the model also maintains successive peaking of increasingly larger tau oligomers, with monomers peaking first and tangles eventually becoming the dominant tau species. This drop-off and plateau of smaller oligomers reflect Smoluchowski aggregation, causing the removal of these oligomers to generate tangles, which increase throughout the time range. However, monomers show a significantly larger peak than other oligomers, primarily due to monomer generation via cytokine-facilitated tau misfolding.

### Proposed model as a Computational Testbed

The model provides a computational testbed for generating mechanistic hypotheses regarding rates difficult to isolate in vivo (Figure 4). We focused on how microglia-dependent tau propagation and pro-inflammatory cytokine release impact global tau burden, and how microglial activation via tau tangles modulates this effect. These parameters were selected for mechanistic importance, as they encompass the neuroinflammatory feedback loop on tau: tau activates microglia, microglia release cytokines causing tau misfolding, microglia contribute to tau propagation, causing more tau aggregation and propagation, restarting the cycle. Microglial activation rate was set to low (0.1x original value), medium (original value), or high (10x original value). We expected a non-linear, yet robust, relation between rate constant scaling and global tau load.

**FIG 4:**
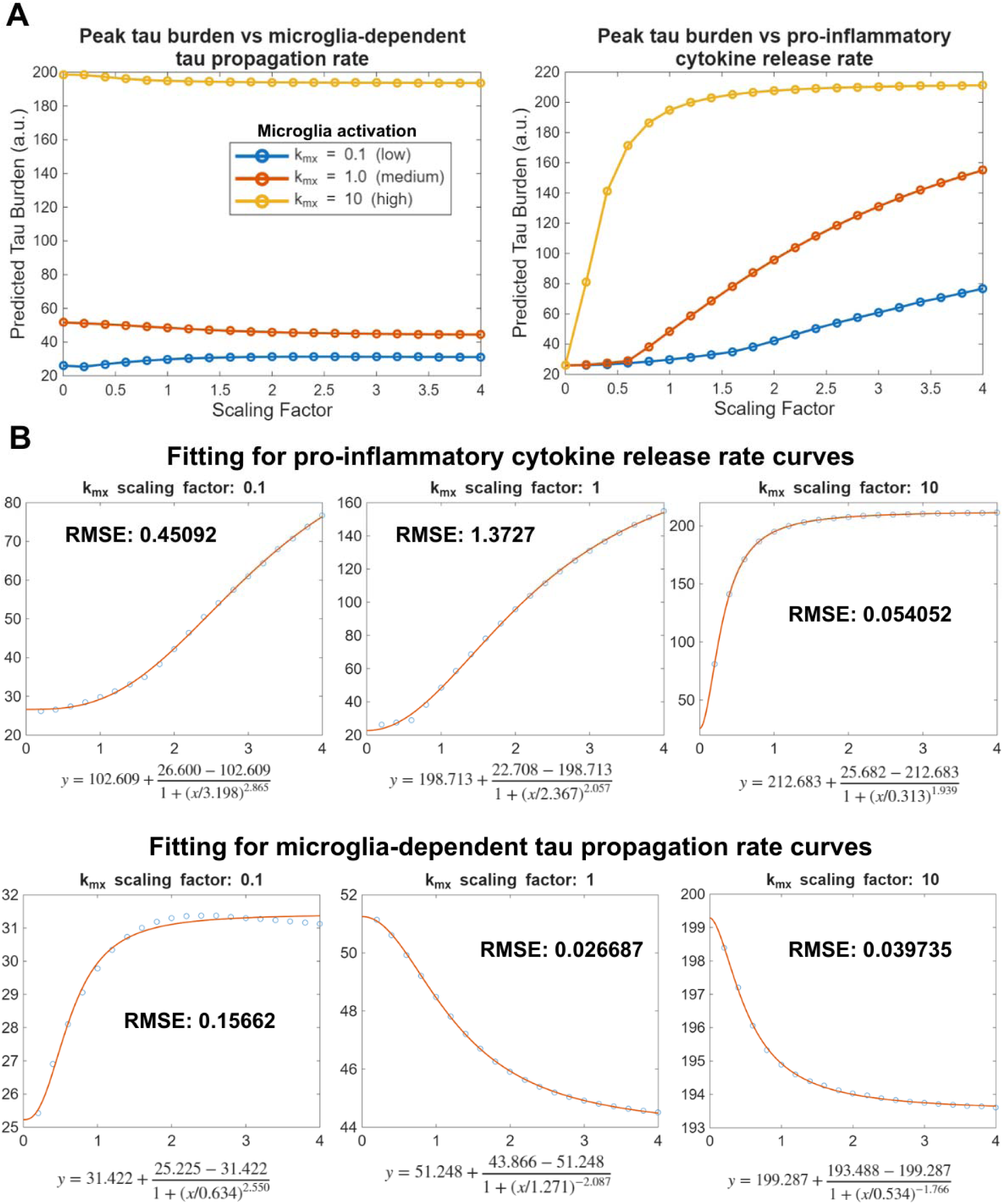
(A) Maximum tau burden plotted over varying rates of microglia-facilitated tau spread (left) and pro-inflammatory cytokine release (right), ranging from a scaling factor of 0.2 to 4.0. This scaling factor was multiplied by the respective parameter’s nominal value. (multiplied by the original parameter value). Additionally, the plot shows variation of three levels of microglial activation (governed by parameter k_mx_ ): low (scaling factor of 0.1), medium (scaling factor of 1, original value) or high (scaling factor of 10). **(B)** Visualization of the fits of a four-parameter sigmoidal function (orange curves) to the relationship between parameter scaling and maximal tau burden (blue dots). Fits were performed separately for each microglial activation level (low, medium, high) and for both the pro-inflammatory cytokine release rate (top) and the microglia-dependent tau propagation rate (bottom). Root mean square error (RMSE) values are also included to assess the quality of the sigmoid fits.

From Figure 4, the relation between parameters and global tau burden is non-linear, following a sigmoidal pattern. These results are biologically plausible, as protein binding rates under cooperative binding typically follow sigmoidal behavior with rapid change at a threshold. The graphs were fitted to a 4-point sigmoidal function (Panel B), which justified a sigmoidal fit because of a low error (measured by root mean square error, RMSE) for all curves.

### Spatiotemporal Tau Trends and Validation

Figure 5 shows predicted regional tau loads compared with experimental progression from regional Flortaucipir-PET scans. For comparison, increasing model time points were chosen to compare with PET scan three subdivisions (naMCI, aMCI, and AD). These time points do not match one-to-one with subdivisions and were chosen to mimic the relative increase in tau distribution (represented by dot size within the glassbrain) as observed in experimental data. Visually, the predicted model fits expected regional progression, with spread from the entorhinal cortex seeding region (green dots in Panel A) to subcortical structures like the hippocampus (black dots), and finally toward parietal and posterior brain regions (blue and purple dots). The most significant glassbrain discrepancy is tau aggregation in the occipital cortex (red dots); otherwise, regional distribution closely mimics increasing PET scan disease severity from naMCI to aMCI to AD.

**FIG 5:**
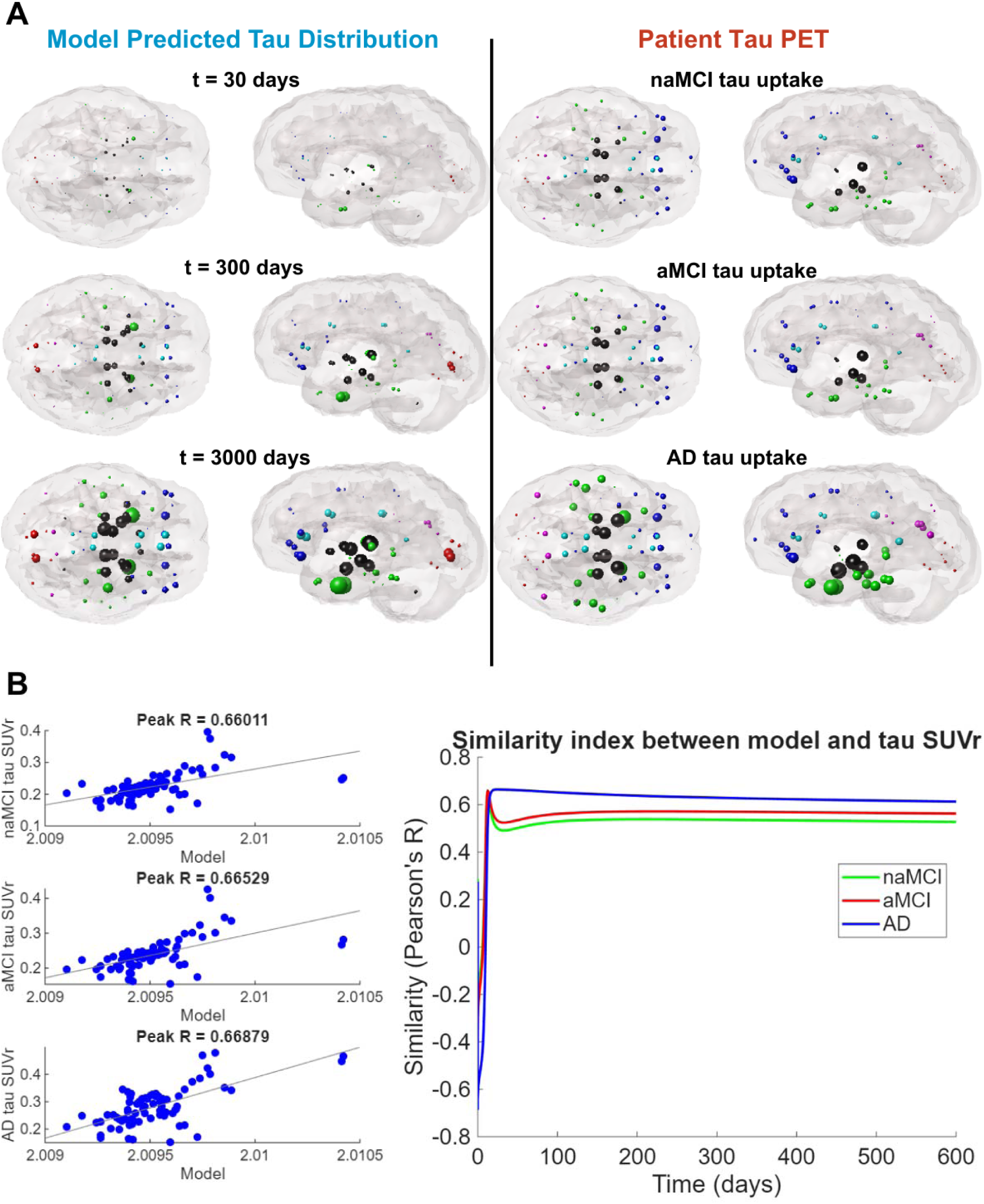
(A) Glassbrain visualization of the model’s predicted tau distribution in parallel to empirical data obtained by regional Florataucipir-PET scans of patients. Color of the spheres represents varying brain regions (blue is frontal, purple is parietal, red is occipital, green is temporal, black is subcortical, and cyan is cingulate). **(B)** Peak Pearson’s R-value and corresponding scatterplots, with the axes representing tau values. The rightmost dots (most amount of tau) represent the entorhinal cortex (EC) which was the seeding region for tau monomers. **(C)** Pearson’s R for the three patient subdivisions over time (600 days), showing the steady-state behavior of the three correlations (naMCI is green, aMCI is red, AD is blue).

For model fit, all three PET data stages show moderately strong correlation (around R=0.66), with slightly better performance for AD data. Interestingly, unlike the AND predecessor which showed markedly better aMCI fit (especially versus naMCI), this model shows consistent Pearson correlations. Finally, R-values for the model and three patient subdivisions were plotted over time (Panel C). AD cohorts displayed quick R increases that held steadily across the time range. For naMCI and aMCI cohorts, R-values peak early then dip slightly before gradually increasing and reaching steady state toward the time range end. This steady-state behavior reflects Pearson correlation’s scale insensitivity, so R-values remain constant as tau increases.

### Spatiotemporal Profile of Inflammation-Related Agents

Similar to glassbrain tau renderings, the final loads of key inflammatory agents – including astrocytes, cytokines, and microglia – were visualized (Figure 6). These agents were modulated with the same Laplacian matrix as tau, but with unique constants governing spread propensity. Despite this, significant differences emerged in pro-inflammatory and anti-inflammatory marker dispersion. Pro-inflammatory markers (M1 microglia, pro-inflammatory cytokines, and astrocytes) exhibited spread toward posterior brain portions, with notable presence in non-temporal/limbic regions. Conversely, anti-inflammatory markers (M2 microglia and anti-inflammatory cytokines) display preferential spread toward anterior regions, including prominent frontal lobe presence, and remain more concentrated in temporal regions (where initial seeding occurred).

**FIG 6:**
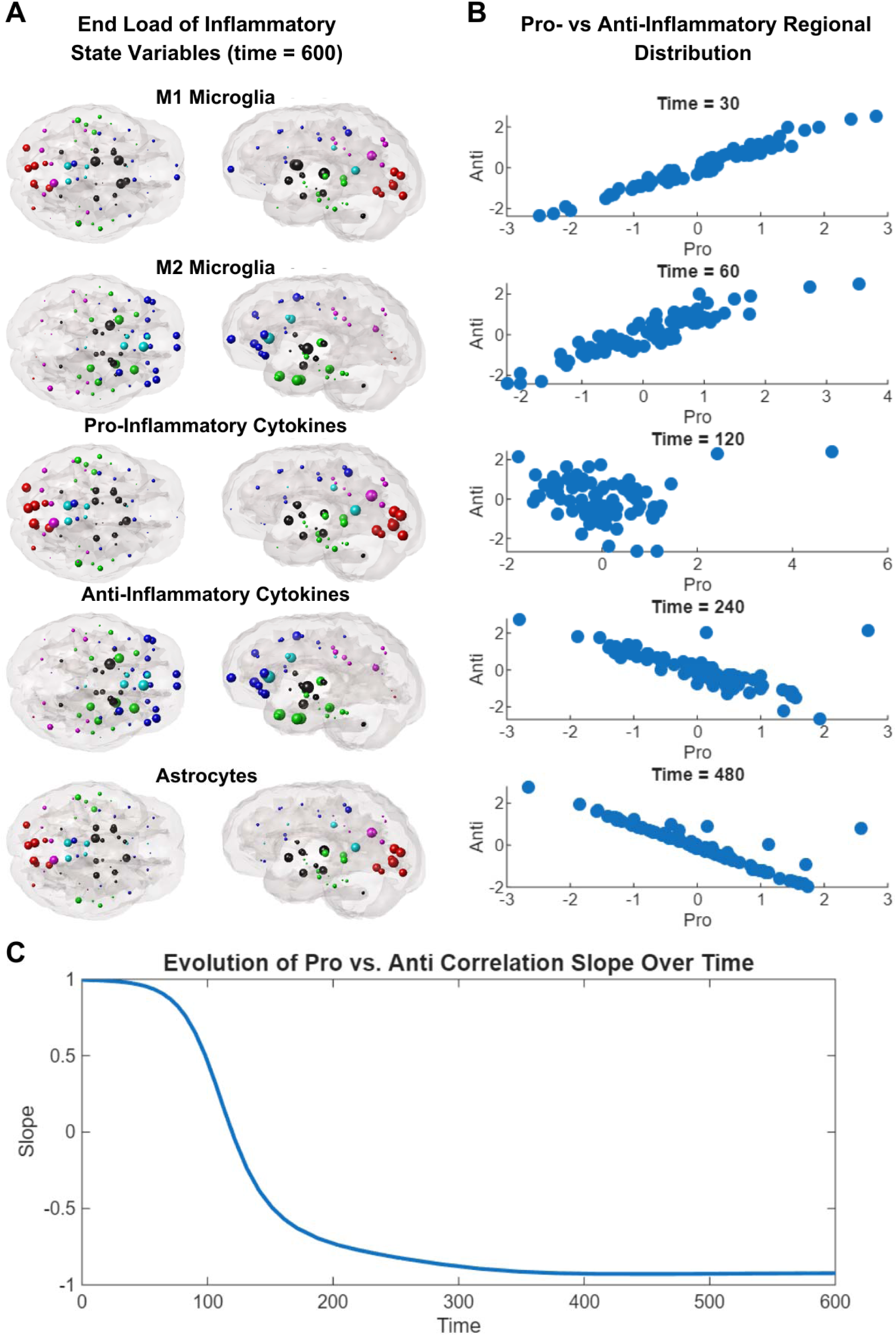
(A) Glassbrain visualization of the regional distribution of M1 microglia, M2 microglia, pro-inflammatory cytokines, anti-inflammatory cytokines, and astrocytes after t=600 days. As for the tau renderings, the different color spheres reflect different brain regions. **(B)** Correlation between the regional distribution of pro- and anti-inflammatory elements at progressing time points, calculated using the average of the z-scores of M1 microglia and pro-inflammatory cytokines, and the same for M2 microglia and anti-inflammatory cytokines. A positive association represents the presence of both pro- and anti- elements in the same region, while a negative association represents an opposing distribution of pro- and anti- elements. **(C)** Plot of the slope of correlations from **(B)** over time. Graph shows a sigmoidal pattern and shows the shift from a positive to a negative association between pro- and anti-inflammatory elements’ regional distribution.

This preferential distribution develops over time rather than being initially modeled, which makes sense considering no initial differences in microglia or cytokine types were incorporated. Panel B visualizes this shift, showing scatter plots of normalized pro-versus anti-inflammatory agent values at different time points. The key metric is slope, which was plotted over time (Panel C), showing whether pro- and anti-inflammatory elements occupy similar brain regions (positive slope), have no regional distribution association (zero slope), or occupy opposing brain regions (negative slope). Around t=120 days, the model shifts from positive to zero slope, showing no systematic association between pro- and anti-inflammatory elements. This pattern continues until, by the time span end, the model shows reversed initial positive regional correlation, with pro- and anti-inflammatory elements appearing in opposing brain regions, producing negative slopes.

### Comparison with null models using randomized gene expression

To explore the role of regional genetic matrices used to incorporate gene-specific effects, we replaced region-specific matrices with randomized ones that scrambled regional expression of both original matrices (one for ApoE and one for TREM2). Using 2000 iterations, Figure 7 shows mean peak R values for three patient groups (naMCI, aMCI, and AD).

**FIG 7:**
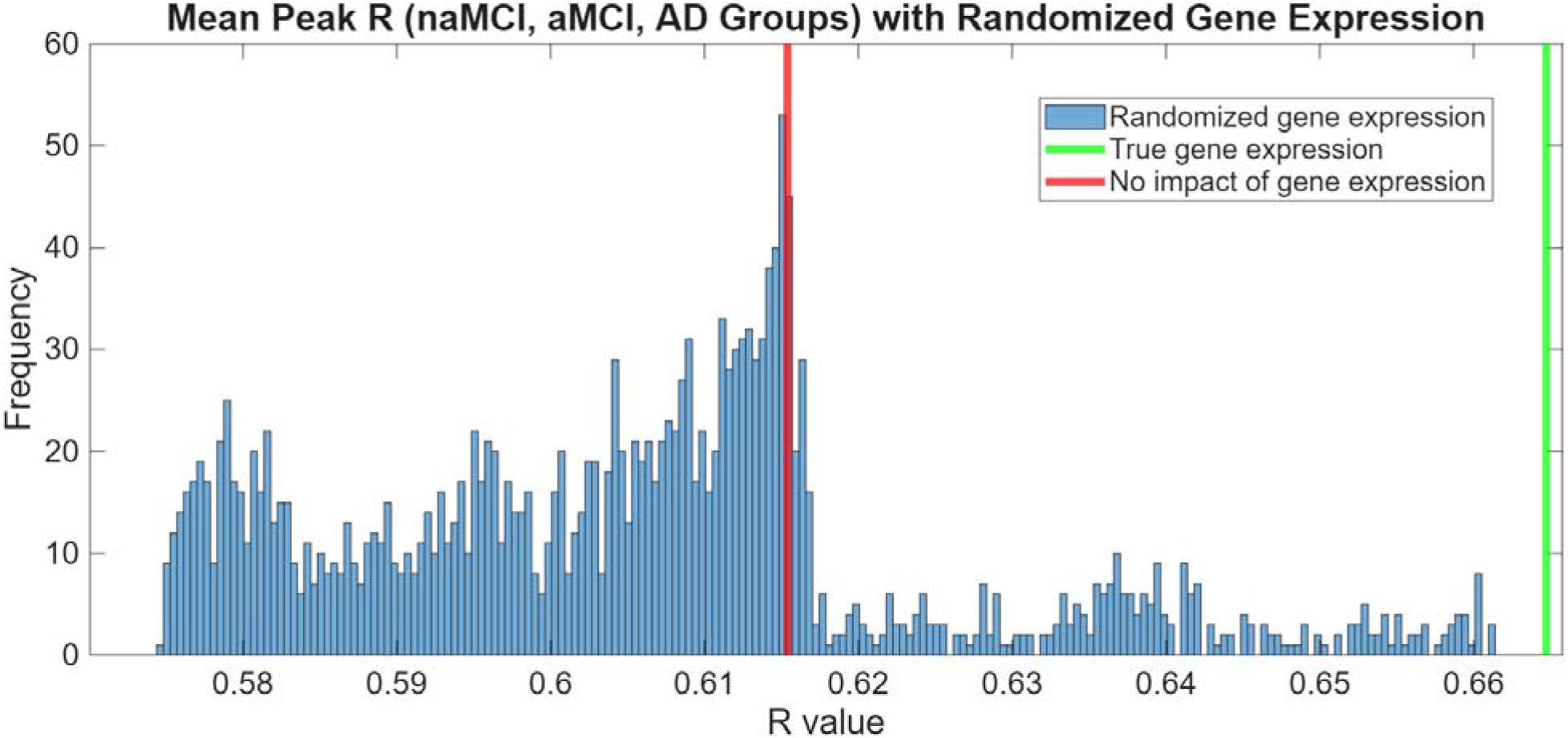
Histogram of mean peak R achieved by model and naMCI, aMCI, and AD cohorts with 2000 trials of gene expression randomly scrambled across brain regions. The trials randomized the two genetic matrices used for TREM2 and ApoE, and this analysis was done on the baseline configuration of wild-type TREM2 and ApoE3. The R-values for the true gene expression (green line) and genetic expression with no regional variance (red line) are shown for comparison.

The model’s original performance (green line) surpasses randomized matrices, with none of 2000 trials producing better mean R. The distribution is roughly unimodal, with the R-value for the model with no region-dependent gene variation (red line) corresponding to the 2000 trial peak. This randomized distribution serves as the null hypothesis for the gene-based model portion, showing expected R values without true, underlying genetic structure or association. Because genetic matrix addition is significant (as no trials surpassed it), this analysis confirms that gene patterns contain inherent structure and demonstrates the non-random role genetic modulation exerts on tau dynamics.

### Regression analysis to assess parameter sensitivity

To understand inflammatory rate parameter impacts, we used linear regression analysis, determining which parameters impact the temporal progression of state variables (focusing on Tau, M1, and Y1) visualized as a heatmap (Figure 8). Rate parameters were varied randomly for 1000 iterations, and Partial Least Squares (PLS) regression characterized parameter impacts on global values of peak tau, ending healthy neuron population, peak M1 microglia, and peak pro-inflammatory cytokines, plus time to reach 50% maximum for M1 and Y1. The heatmap showed several parameters had little impact on these outputs, instead mediating state variables not explicitly measured (like astrocytes or TGF-beta) that are mechanistically helpful but less important for tau dynamics.

**FIG 8:**
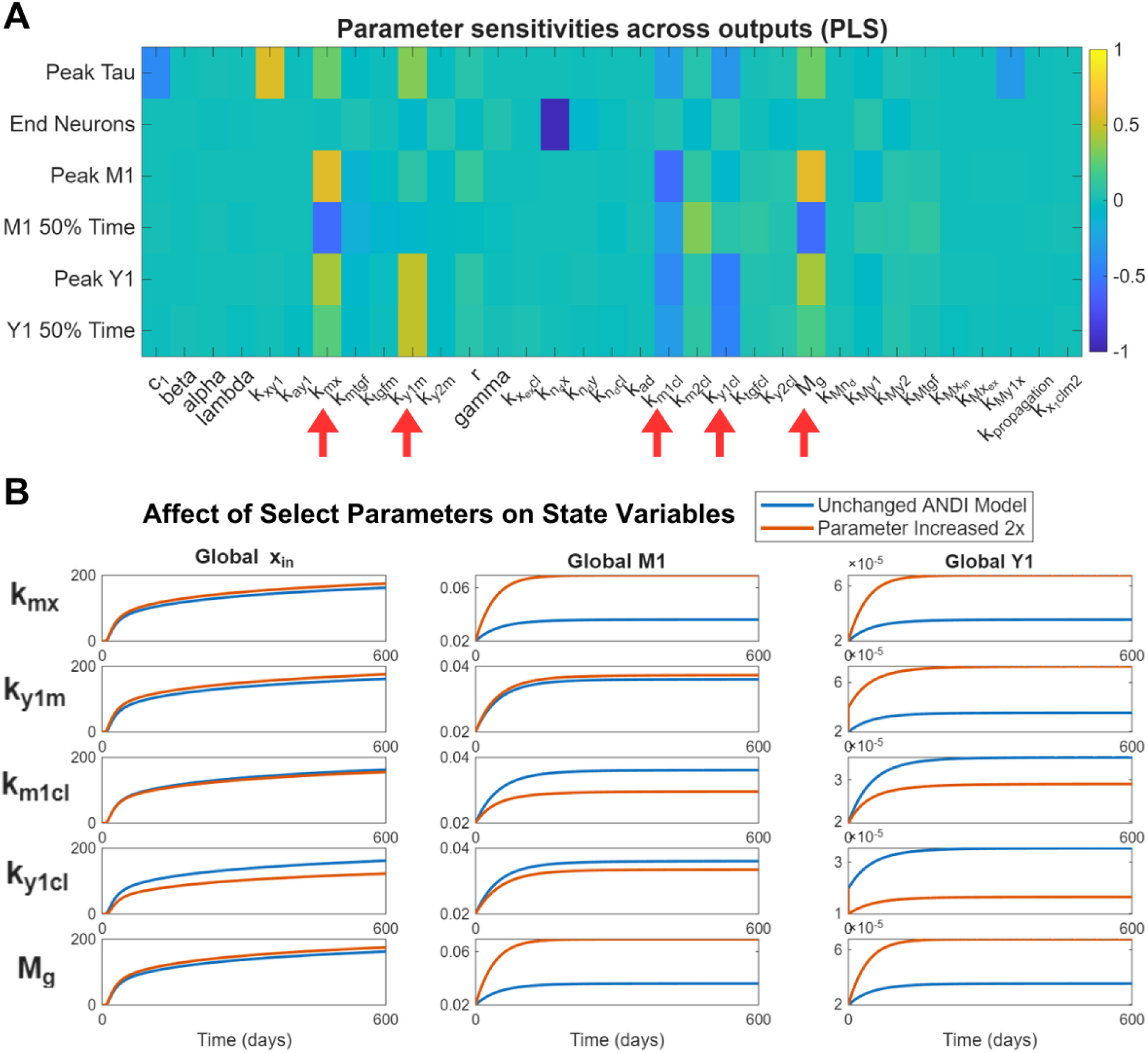
(A) Heatmap visualization of parameter sensitivity on outputs relating to key state variables of tau, healthy neurons, M1 microglia, and pro-inflammatory cytokines, with a dark blue suggesting the parameter decreases an output’s value and brighter yellow suggesting the parameter increases an output’s value. Here, PLS (partial least squares) regression was run on a set of 1000 randomized parameter variations. **(B)** Impact of the five most influential parameters from **(A)** on global temporal evolution of tau, M1 microglia, and pro-inflammatory microglia. The graph shows the impact of increasing each of the selected parameters by a multiplicative factor of 2 (orange) and comparing it with the unchanged ANDI model (blue).

Hence five key parameters were selected based on prominent effect size: microglia activation rate by tau ( k_mx_ ) pro-inflammatory cytokine release rate by M1 microglia (k_ylm_ ), M1 microglia clearance rate (k_mlcl_ ), pro-inflammatory cytokine clearance rate ( k_ylcl_ ), and microglia source for M1 or M2 activation ( M_g_ ). Panel B visualizes how increasing these selected parameters impacts global tau, M1, and Y1 values. Increases in k_mx_ and M_g_primarily cause an increase in M1 and Y1, with smaller (and indirect) tau effects. Increasing k_ylm_ , has similar pro-inflammatory impacts, but primarily on Y1. Finally, as expected, increasing the selected clearance variables (k_ylcl_, k_mlcl_) is anti-inflammatory and neuroprotective, decreasing tau, M1, and Y1.

### Inflammatory Module Contribution to Model Behavior and Pearson’s R-value

Figure 9 compares the baseline model to one with the inflammatory module removed to better understand inflammation’s effect on spatiotemporal tau propagation. First, global tau dynamics were compared (Panel A). The model without inflammation displayed lower tau species values, attributable to cytokine-facilitated monomer generation. However, beyond magnitude, the progression of tau varies significantly. By normalizing the model such that the peak tau of both the ANDI and base AND model were equivalent, we compared the shape and overall progression of tau dynamics. First, looking at monomers and small oligomers (m=2), the model with inflammation displays a greater presence of tau later into the time span, suggesting the added inflammatory processes promote the generation smaller tau aggregates. For larger oligomers (m=4), the model without inflammation displays a wider and more sustained peak, and for tangles, the model without inflammation is much quicker to reach a peak value. Taken together, these differences suggest the added inflammatory processes promote the generation of tau monomers and oppose the aggregation of larger oligomers, which culminates in a slower and drawn-out evolution of tangles.

**FIG 9:**
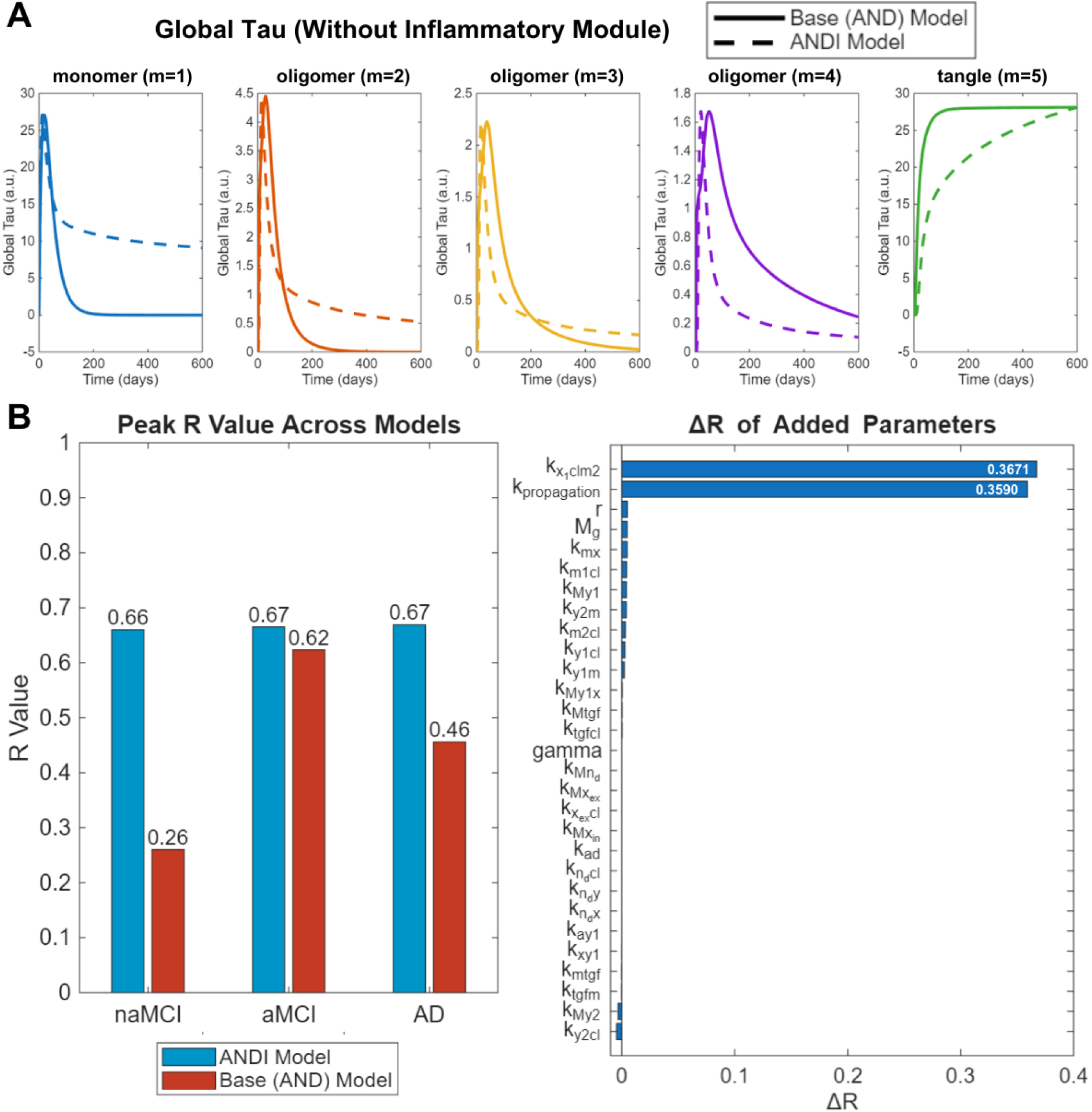
(A) Comparison of the base (AND) model without inflammation and the ANDI model on global tau of varying lengths. To facilitate comparison of the shape and behavior of the two, ANDI model behavior was scaled via max normalization, such that the peak tau values of it would be equivalent to the model without inflammation. **(B)** Comparison of the peak Pearson’s R across the three patient subdivisions (naMCI, aMCI, AD) between the no-inflammation base (AND) and the ANDI model. On the left, visualization of each rate parameter’s contribution towards the increased fit. This parameter contribution was computed by taking a delta R, subtracting the similarity (Pearson’s R) of the complete model and the ANDI model in absence of that parameter.

We also computed this non-inflammatory model’s correlation across the same time span (Panel B, right). Compared to the ANDI model, the base no-inflammation model shows markedly lower peak R values across all three patient subdivisions. The most prominent difference is in the naMCI cohort (0.6601 vs. 0.2604). The AD cohort shows a smaller but still marked difference (0.6688 vs. 0.4560), and the aMCI cohort has similar peak R across both models (0.6653 vs. 0.6233).

We then characterized various inflammation-related rate contributions to this R increase (Panel B, left). ΔR was computed, subtracting the model’s mean peak R-value for each of the three patient groups from the corresponding R when each rate constant has no effect. Most parameters contribute positively toward R-values, with several having negligible impact. The most prominent rate constants were rates at which M1 microglia spread tau ( ), and M2 microglia clear tau ( ), with ΔRs of 0.3590 and 0.3671 respectively. These rate constants directly modify tau values, as opposed to indirect or downstream effects. Interestingly, both parameters involve microglia, indicating higher microglia importance versus other state variables for model predictive ability.

### Creating a More Parsimonious Model

Figure 9 characterizes model parameters essential for model fit, allowing creation of a minimal, more parsimonious model (Figure 10). This model was created by removing parameters (by setting to 0) that did not contribute to similarity with experimental tau SUVr data, while maintaining overall behavior, trends, and stability. Subsequently, several state variables were made constants depending on whether their rate constants were removed. This streamlining left tau-related state variables and only M1 and M2 microglia for inflammation-related state variables.

**FIG 10:**
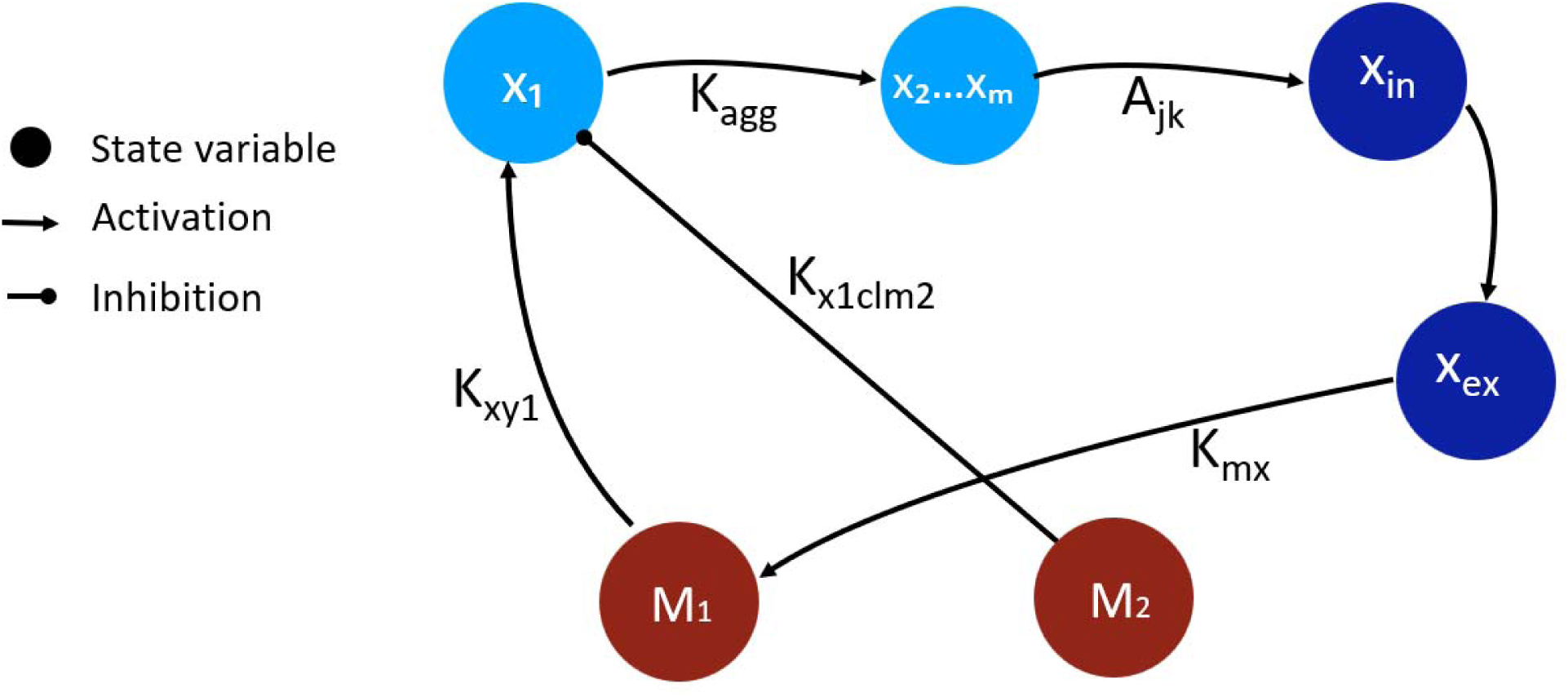
Schematic showing the remaining interactions in the parsimonious model. Again, clearance rates and parameters impacting propagation are not included in the diagram. Of the original state variables, only tau-related variables and M1 and M2 microglia remain.

To assess the minimal model, we then compared it to the complete ANDI model, judging the model’s ability to maintain biologically plausible state variable progressions and similarity (Pearson’s R) with empirical tau data (Figure 11). Panel A compares time-dependent evolution of the similarity index for naMCI, aMCI, and AD between the complete ANDI model and the minimal model, showing nearly identical peak Pearson’s R and temporal progression of the R-value across all three subdivisions (naMCI, aMCI, AD). This confirms that the parsimonious model retains value for spatiotemporal tau dynamics, despite lacking some mechanistic relations. Panel B shows global behavior for tau, M1 microglia, and M2 microglia in both models, showing that the minimal model displays higher global tau and M1 as well as lower global M2. Because of these discrepancies in state variable progression (favoring M1, opposing M2 in minimal model), the dynamics kept in the minimal model seem to be more oriented towards the pro-inflammatory cascade.

**FIG 11:**
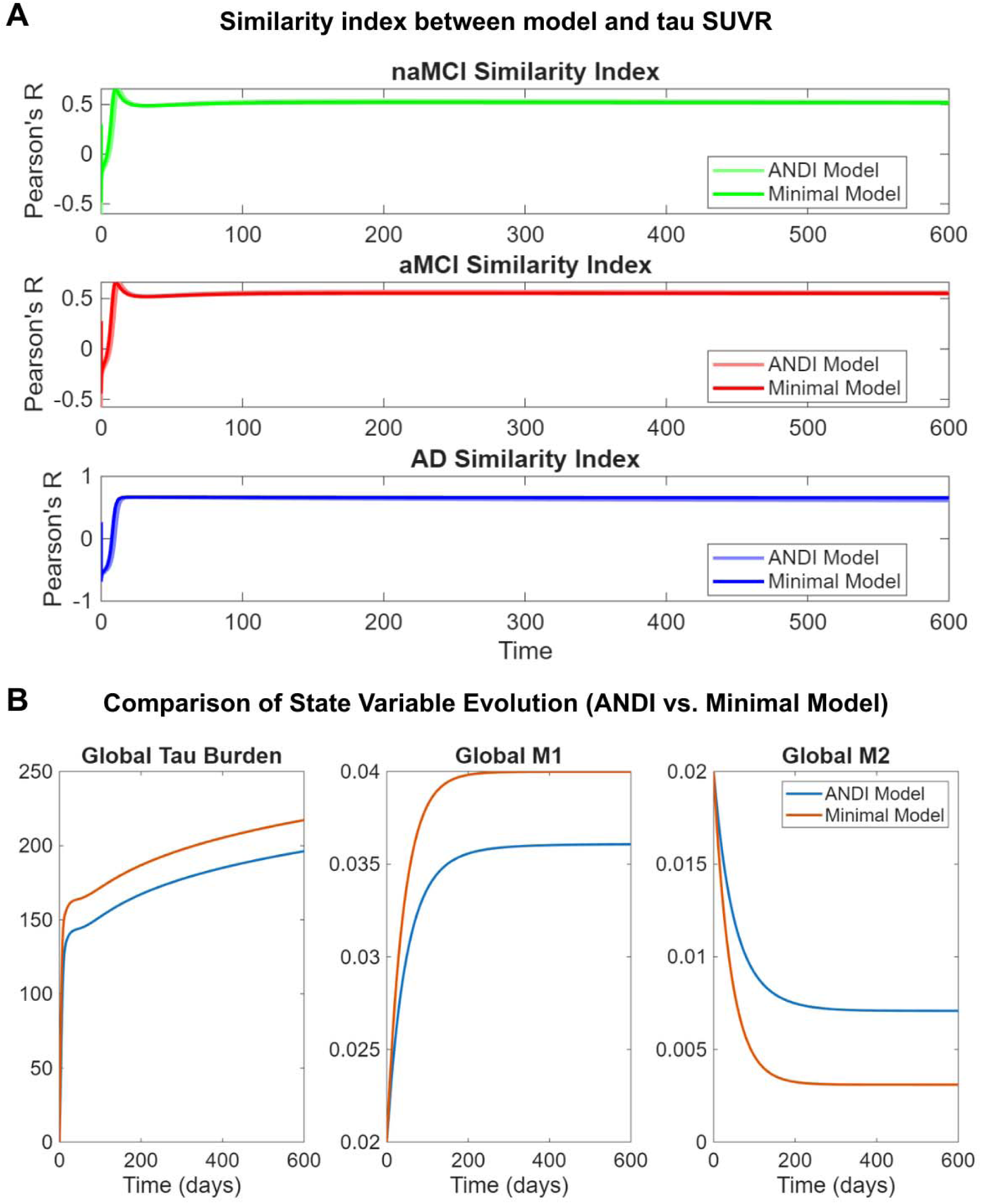
(A) Comparison of similarity to Tau SUVr with the minimal model (darker colors) and the ANDI model (lighter color) for all three patient subdivisions (naMCI, aMCI, AD). **(B)** Comparison of the minimum model (orange) and the ANDI model (blue) in the global evolution of the remaining state variables (tau, M1 microglia, and M2 microglia) over a time range of 600 days.

## Discussion

This work presents an extended computational framework integrating tau propagation with neuroinflammatory and genetic pathways, successfully augmenting model similarity for regional tau distribution across Alzheimer’s progression stages (Figure 8). We built upon the AND model’s framework, chosen for its simple addition of inflammatory dynamics in rigorous examination of tau propagation across specific brain regions. Within each local brain region, we incorporated key inflammatory mechanisms including microglial activation, cytokine release, neuronal death, and genetic impacts for ApoE and TREM2, examining how these processes affected brain-wide tau spread. The major findings demonstrate neuroinflammation’s and related genetic impacts’ computational value for modeling tau propagation in AD.

### Computational value of neuroinflammation and genetic pathways

A central outcome of the presented framework is its recapitulation of the role of inflammatory processes, by demonstrating that the added inflammatory module significantly increased model similarity (Pearson’s R) evaluated on regional tau SUVr. Unlike prior tau-only implementations (like the base AND model), the presented ANDI model maintains very similar indices over all three patient subdivisions (naMCI, aMCI, AD) with R values around 0.66, departing from the disproportionally higher aMCI fit with far poorer naMCI fit seen in the base model. Because naMCI cases tend to be heterogeneous, the traditional tau pathology seen in AD is not necessarily a major driver. Thus, ANDI’s increased similarity can be attributed to neuroinflammation’s more universal role in cognitive impairment, and its role as a major driver of AD onset (48, 49). This preclinical benefit neuroinflammation also correlates with experimental studies suggesting anti-inflammatory agents may act as important protective factors in presymptomatic intervention (50).

Regarding genetic pathways, the model demonstrated that gene data provides R-value improvement across subdivisions, with higher similarity to tau SUVr than 2000 randomly scrambled genetic matrices. These results show potential significance for the selective vulnerability hypothesis, suggesting predisposed genetic distribution of TREM2 and ApoE influences regional vulnerability in AD, although the fact that genetic expression caused a relatively small increase in model R suggests it is not the sole factor behind regional spread of tau. Taken together with the aforementioned role of inflammatory dynamics, the ANDI model provides a powerful argument for the incorporation of neuroinflammation and genetic pathways in modeling efforts, especially considering their importance in capturing regional variance that are inadequately explained by tau network spread alone.

This computational value seems to, in large part, stem from the incorporation of microglia. By leveraging the model’s role as a computational testbed, we explored hypotheses centered on microglia-related parameters, visualizing global tau behavior as a function of microglia-mediated tau propagation and pro-inflammatory cytokine release (Figure 3). As expected, inflammatory modulations of tau aggregation, like cytokine release rates, displayed clear relations with global tau burden; however, this tau burden impact was also seen in inflammatory modulations of tau propagation (through microglial tau spread rates), despite global tau not being as related by regional spread, supporting the central role of microglia in shaping tau dynamics. For microglia-mediated propagation specifically, lower microglial activation rates show a positive relationship between microglial spread rate and global tau burden, while at higher microglial activation this relation is negative. This furthers the notion that microglia play larger roles in progressing tau pathology during early AD, when microglia activation rates are lower.

### Emergent pro- and anti-inflammatory spatial divergence

One of the most notable behaviors to emerge from the model is the progressive spatial divergence between pro- and anti-inflammatory elements (Figure 5). Despite identical initial distributions and graph diffusion term of microglia and cytokine subtypes, the model produces a time-dependent segregation, with pro-inflammatory markers preferentially occupying posterior regions of the brain and anti-inflammatory markers remaining concentrated in anterior areas. This divergence may be attributed to phenomena of two inflammatory peaks (51), such that anti-inflammatory markers peak in regions implicated early in AD and pro-inflammatory correspond to the secondary peak and thus affect later-stage regions (like the parietal lobe). Because the pro- and anti- components were subject to the same modeling constraints, this divergence also suggests that network transmissibility suffices in explaining regional vulnerability and generating a heterogenous inflammatory landscape. Combined with conclusions from genetic regional variance in the above section, the model shows a more dynamic and nuanced view of regional vulnerability with network transmission as a dominant driver behind spatial patterns, modulated by predisposed factors like gene expression.

### Model reduction and parsimony

Given the inherent and significant challenges of parameter identifiability in models with high dimensionality, the identification of a more parsimonious model retaining only tau and microglial state variables is key and informative (Figure 9). Despite the removal of multiple state variables and rate constants, the minimal model preserves nearly identical similarity to regional tau PET data, with only tau, M1 microglia, and M2 microglia as time-dependent state variables and other variables as constants. These findings lay the groundwork for future work. In modeling neuroinflammation in AD, several rate constants and parameters cannot be accurately quantified due to limited experimental support, and our results show we can significantly reduce model uncertainty (via fewer state variables) while maintaining robustness, at least for tau propagation.

Regarding parameters, we identified microglia-mediated tau spread and M2 microglia tau clearance rates as major contributors toward accurate tau propagation representation. This mirrors developing experimental understanding of microglial modulation of tau spread (52) and simultaneous clearance effects (53); however, the extent to which these two parameters define model fit is intriguing, with both representing about 0.36 R increases, a stark difference from all other parameters with delta R under 0.01 (Figure 9). The minimal model also showed pro-inflammatory cascades are more important to model for tau spread, evidenced by its transformation of state variable progression towards increased tau and M1 microglia, and decreased M2 microglia concentrations. Thus, our work shows that for creating efficient and parsimonious models of neuroinflammation and tau, modeling efforts should prioritize microglia as time-dependent variables, with microglia-dependent tau spread and degradation rate constants, and emphasis on more explicit pro-inflammatory cascade modeling.

### Note on amyloid beta

AD research has historically focused on amyloid pathology, dominated by the amyloid cascade hypothesis positing that upstream Aβ accumulation triggers tau pathology and downstream neurodegeneration. However, as mentioned before, most Aβ-targeting drugs have failed to significantly alter AD pathology (35). Thus, rather than explicitly modeling amyloid dynamics, the presented framework treats inflammation as an intermediary through which amyloid-related effects may influence tau aggregation and propagation. To that point, recent modeling efforts suggest that both tau and Aβ employ network-mediated spread (18, 54, 55). This scheme allows complex interactions between species, supported experimentally by amyloid precursor protein (APP) promoting tau aggregation (56), tau influencing Aβ formation (57), and the AD-specific retrograde spread of tau potentially being due to modulation via Aβ (58). Importantly, neuroinflammation exists as an intermediary between bidirectional tau and Aβ effects, with studies proposing Aβ modulates tau propagation and aggregation via pro-inflammatory microglial responses (52, 59).

### Potential extensions and broader applications

Our model focused on tau propagation, and neuroinflammation as a secondary modulator. However, our parsimonious model identification could, through its simplicity, facilitate more rigorously validated models and analyses of regional neuroinflammation spread. Because a major hurdle is lack of parameter identifiability from available data, this simpler yet biologically accurate model could reduce required data and ease construction viability. More broadly, given tau and inflammation universality in various disorders (including Parkinson’s, frontotemporal dementia, progressive supranuclear palsy, corticobasal degeneration, amyotrophic lateral sclerosis, etc.), our model presents a viable basis for understanding driving pathologies. Inflammatory dynamics are largely consistent across these disorders; with necessary data (via CSF biomarkers, for example), this model can extend to reflect disease-specific dynamics. Finally, the model can help pinpoint underlying processes behind genetic impacts in AD. For example, with regional gene expression data (like our ApoE and TREM2 implementations), sophisticated quantification of how gene expressions impact state variables could allow researchers to work backwards and identify rate parameters bridging gene expression inputs and pathological outputs. Beyond insights from analysis of model behavior, our work can serve as a computational testbed, bridging hypothetical and theoretical science with more concrete experimental imaging studies without intensive resource demands. We believe this work can catalyze further and more detailed examinations of neuroinflammation and tau interplay in neurodegenerative diseases.

### Limitations

This study has several limitations. The most significant is the high dimensionality of the parameter space. The model contains many biological rate constants, many of which cannot be directly inferred from available human or animal data. Our results could thus be subject to substantial parameter non-identifiability, meaning multiple parameter value combinations could reproduce similar disease trajectories and limit derivable insights. We mitigated this through a more parsimonious model; nevertheless, further work is needed to address this lack of rigorous basis for extensive parameters needed for mechanistic modeling. Also, the cytokine module was simplified into pro- and anti-inflammatory categories for computational simplicity, but this omits diverse signaling cascade diversity (including IL-6, chemokine families, etc.) known to shape microglial states diversely and combined distinct effects to better recreate inflammatory responses. Similarly, M1/M2 microglia duality was a simplification, ignoring mounting evidence that microglia exhibit more nuanced states (60). Additionally, using static parameters across individuals and regions neglects patient- and region-specific variability in tau kinetics and immune responsiveness. These simplifications limit immediate clinical applicability. However, overall result consistency suggests inflammation and genetics inclusion helps capture AD progression features unexplained by tau propagation alone.

Another limitation concerns temporal scale. Though global inflammatory trends use non-arbitrary time scales (days), reflected in parameter values, the model lacks temporal utility. For example, though the time variable reflects accurate global trends, it should not be interpreted as direct mapping to clinical disease stages from naMCI to aMCI to AD. This is because Pearson’s correlation, our model performance metric, does not factor in scale, instead evaluating how well the model captures spatial patterns of tau distribution across regions, without accounting for absolute tau burden. The model optimizes reproduction of propagation patterns, with plausible, but not necessarily quantitative, tau aggregation measures. This, though a significant limitation, reflects our study focus, evaluating insights on how regional tau spread and propagation are modulated by neuroinflammation and genetics, and this modulation’s significance on model performance.

## References

1. Selkoe DJ, Lansbury PJ Jr. Alzheimer’s Disease Is the Most Common Neurodegenerative Disorder. In: Siegel GJ, Agranoff BW, Albers RW, et al., editors. Basic Neurochemistry: Molecular, Cellular and Medical Aspects. 6th edition. Philadelphia: Lippincott-Raven; 1999. Available from: https://www.ncbi.nlm.nih.gov/books/NBK27944/

2. 2024 Alzheimer’s disease facts and figures. Alzheimers Dement. 2024 May;20(5):3708–3821. doi: 10.1002/alz.13809. Epub 2024 Apr 30. PMID: 38689398; PMCID: PMC11095490.

3. Masters CL, Simms G, Weinman NA, Multhaup G, McDonald BL, Beyreuther K. Amyloid plaque core protein in Alzheimer disease and Down syndrome. Proc Natl Acad Sci U S A. 1985 Jun;82(12):4245–9. doi: 10.1073/pnas.82.12.4245. PMID: 3159021; PMCID: PMC397973.

4. KIDD M. Paired helical filaments in electron microscopy of Alzheimer’s disease. Nature. 1963 Jan 12;197:192–3. doi: 10.1038/197192b0. PMID: 14032480.

5. Heneka MT, Carson MJ, El Khoury J, Landreth GE, Brosseron F, Feinstein DL, Jacobs AH, Wyss-Coray T, Vitorica J, Ransohoff RM, Herrup K, Frautschy SA, Finsen B, Brown GC, Verkhratsky A, Yamanaka K, Koistinaho J, Latz E, Halle A, Petzold GC, Town T, Morgan D, Shinohara ML, Perry VH, Holmes C, Bazan NG, Brooks DJ, Hunot S, Joseph B, Deigendesch N, Garaschuk O, Boddeke E, Dinarello CA, Breitner JC, Cole GM, Golenbock DT, Kummer MP. Neuroinflammation in Alzheimer’s disease. Lancet Neurol. 2015 Apr;14(4):388–405. doi: 10.1016/S1474-4422(15)70016-5. PMID: 25792098; PMCID: PMC5909703.

6. Chen Y, Yu Y. Tau and neuroinflammation in Alzheimer’s disease: interplay mechanisms and clinical translation. J Neuroinflammation. 2023 Jul 14;20(1):165. doi: 10.1186/s12974-023-02853-3. PMID: 37452321; PMCID: PMC10349496.

7. Weingarten MD, Lockwood AH, Hwo SY, Kirschner MW. A protein factor essential for microtubule assembly. Proc Natl Acad Sci U S A. 1975 May;72(5):1858–62. doi: 10.1073/pnas.72.5.1858. PMID: 1057175; PMCID: PMC432646.

8. Merrick SE, Demoise DC, Lee VM. Site-specific dephosphorylation of tau protein at Ser202/Thr205 in response to microtubule depolymerization in cultured human neurons involves protein phosphatase 2A. J Biol Chem. 1996 Mar 8;271(10):5589–94. doi: 10.1074/jbc.271.10.5589. PMID: 8621419.

9. Frost B, Diamond MI. Prion-like mechanisms in neurodegenerative diseases. Nat Rev Neurosci. 2010 Mar;11(3):155–9. doi: 10.1038/nrn2786. Epub 2009 Dec 23. PMID: 20029438; PMCID: PMC3648341.

10. Christianson HC, Belting M. Heparan sulfate proteoglycan as a cell-surface endocytosis receptor. Matrix Biol. 2014 Apr;35:51–5. doi: 10.1016/j.matbio.2013.10.004. Epub 2013 Oct 18. PMID: 24145152.

11. Pérez M, Avila J, Hernández F. Propagation of Tau via Extracellular Vesicles. Front Neurosci. 2019 Jul 2;13:698. doi: 10.3389/fnins.2019.00698. PMID: 31312118; PMCID: PMC6614378.

12. Masel J, Jansen VA, Nowak MA. Quantifying the kinetic parameters of prion replication. Biophys Chem. 1999 Mar 29;77(2-3):139–52. doi: 10.1016/s0301-4622(99)00016-2. PMID: 10326247.

13. Prusiner SB, Scott M, Foster D, Pan KM, Groth D, Mirenda C, Torchia M, Yang SL, Serban D, Carlson GA, et al. Transgenetic studies implicate interactions between homologous PrP isoforms in scrapie prion replication. Cell. 1990 Nov 16;63(4):673–86. doi: 10.1016/0092-8674(90)90134-z. PMID: 1977523.

14. Bertsch M, Franchi B, Marcello N, Tesi MC, Tosin A. Alzheimer’s disease: a mathematical model for onset and progression. Math Med Biol. 2017 Jun 1;34(2):193–214. doi: 10.1093/imammb/dqw003. PMID: 27079222.

15. Matthäus F. Diffusion versus network models as descriptions for the spread of prion diseases in the brain. J Theor Biol. 2006 May 7;240(1):104–13. doi: 10.1016/j.jtbi.2005.08.030. Epub 2005 Oct 10. PMID: 16219329.

16. Iturria-Medina Y, Sotero RC, Toussaint PJ, Evans AC; Alzheimer’s Disease Neuroimaging Initiative. Epidemic spreading model to characterize misfolded proteins propagation in aging and associated neurodegenerative disorders. PLoS Comput Biol. 2014 Nov 20;10(11):e1003956. doi: 10.1371/journal.pcbi.1003956. PMID: 25412207; PMCID: PMC4238950.

17. Raj A, Kuceyeski A, Weiner M. A network diffusion model of disease progression in dementia. Neuron. 2012 Mar 22;73(6):1204–15. doi: 10.1016/j.neuron.2011.12.040. Epub 2012 Mar 21. PMID: 22445347; PMCID: PMC3623298.

18. Raj A, Torok J, Ranasinghe K. Understanding the complex interplay between tau, amyloid and the network in the spatiotemporal progression of Alzheimer’s disease. Prog Neurobiol. 2025 Jun;249:102750. doi: 10.1016/j.pneurobio.2025.102750. Epub 2025 Mar 17. PMID: 40107380; PMCID: PMC12542852.

19. Tora V, Torok J, Bertsch M, Raj A. A network-level transport model of tau progression in the Alzheimer’s brain. Math Med Biol. 2025 Mar 17;42(2):212–238. doi: 10.1093/imammb/dqaf003. PMID: 40080630.

20. Dan T, Dere M, Kim WH, Kim M, Wu G. TauFlowNet: Revealing latent propagation mechanism of tau aggregates using deep neural transport equations. Med Image Anal. 2024 Jul;95:103210. doi: 10.1016/j.media.2024.103210. Epub 2024 May 17. PMID: 38776842.

21. Perea JR, López E, Díez-Ballesteros JC, Ávila J, Hernández F, Bolós M. Extracellular Monomeric Tau Is Internalized by Astrocytes. Front Neurosci. 2019 May 1;13:442. doi: 10.3389/fnins.2019.00442. PMID: 31118883; PMCID: PMC6504834.

22. Domingues C, da Cruz E Silva OAB, Henriques AG. Impact of Cytokines and Chemokines on Alzheimer’s Disease Neuropathological Hallmarks. Curr Alzheimer Res. 2017;14(8):870–882. doi: 10.2174/1567205014666170317113606. PMID: 28317487; PMCID: PMC5543563.

23. Dikmen HO, Hemmerich M, Lewen A, Hollnagel JO, Chausse B, Kann O. GM-CSF induces noninflammatory proliferation of microglia and disturbs electrical neuronal network rhythms in situ. J Neuroinflammation. 2020 Aug 11;17(1):235. doi: 10.1186/s12974-020-01903-4. PMID: 32782006; PMCID: PMC7418331.

24. Jope RS, Yuskaitis CJ, Beurel E. Glycogen synthase kinase-3 (GSK3): inflammation, diseases, and therapeutics. Neurochem Res. 2007 Apr-May;32(4-5):577–95. doi: 10.1007/s11064-006-9128-5. Epub 2006 Aug 30. PMID: 16944320; PMCID: PMC1970866.

25. Lee SH, Meilandt WJ, Xie L, Gandham VD, Ngu H, Barck KH, Rezzonico MG, Imperio J, Lalehzadeh G, Huntley MA, Stark KL, Foreman O, Carano RAD, Friedman BA, Sheng M, Easton A, Bohlen CJ, Hansen DV. Trem2 restrains the enhancement of tau accumulation and neurodegeneration by β-amyloid pathology. Neuron. 2021 Apr 21;109(8):1283–1301.e6. doi: 10.1016/j.neuron.2021.02.010. Epub 2021 Mar 5. PMID: 33675684.

26. Liu W, Taso O, Wang R, Bayram S, Graham AC, Garcia-Reitboeck P, Mallach A, Andrews WD, Piers TM, Botia JA, Pocock JM, Cummings DM, Hardy J, Edwards FA, Salih DA. Trem2 promotes anti-inflammatory responses in microglia and is suppressed under pro-inflammatory conditions. Hum Mol Genet. 2020 Nov 25;29(19):3224–3248. doi: 10.1093/hmg/ddaa209. PMID: 32959884; PMCID: PMC7689298.

27. Kwon Y, Mehta S, Clark M, Walters G, Zhong Y, Lee HN, Sunahara RK, Zhang J. Non-canonical β-adrenergic activation of ERK at endosomes. Nature. 2022 Nov;611(7934):173–179. doi: 10.1038/s41586-022-05343-3. Epub 2022 Oct 26. PMID: 36289326; PMCID: PMC10031817.

28. Peng X, Guo H, Zhang X, Yang Z, Ruganzu JB, Yang Z, Wu X, Bi W, Ji S, Yang W. TREM2 Inhibits Tau Hyperphosphorylation and Neuronal Apoptosis via the PI3K/Akt/GSK-3β Signaling Pathway In vivo and In vitro. Mol Neurobiol. 2023 May;60(5):2470–2485. doi: 10.1007/s12035-023-03217-x. Epub 2023 Jan 20. PMID: 36662361.

29. Zhao N, Qiao W, Li F, Ren Y, Zheng J, Martens YA, Wang X, Li L, Liu CC, Chen K, Zhu Y, Ikezu TC, Li Z, Meneses AD, Jin Y, Knight JA, Chen Y, Bastea L, Linares C, Sonustun B, Job L, Smith ML, Xie M, Liu YU, Umpierre AD, Haruwaka K, Quicksall ZS, Storz P, Asmann YW, Wu LJ, Bu G. Elevating microglia TREM2 reduces amyloid seeding and suppresses disease-associated microglia. J Exp Med. 2022 Dec 5;219(12):e20212479. doi: 10.1084/jem.20212479. Epub 2022 Sep 15. PMID: 36107206; PMCID: PMC9481739.

30. Corder EH, Saunders AM, Strittmatter WJ, Schmechel DE, Gaskell PC, Small GW, Roses AD, Haines JL, Pericak-Vance MA. Gene dose of apolipoprotein E type 4 allele and the risk of Alzheimer’s disease in late onset families. Science. 1993 Aug 13;261(5123):921–3. doi: 10.1126/science.8346443. PMID: 8346443.

31. Shi Y, Yamada K, Liddelow SA, Smith ST, Zhao L, Luo W, Tsai RM, Spina S, Grinberg LT, Rojas JC, Gallardo G, Wang K, Roh J, Robinson G, Finn MB, Jiang H, Sullivan PM, Baufeld C, Wood MW, Sutphen C, McCue L, Xiong C, Del-Aguila JL, Morris JC, Cruchaga C; Alzheimer’s Disease Neuroimaging Initiative; Fagan AM, Miller BL, Boxer AL, Seeley WW, Butovsky O, Barres BA, Paul SM, Holtzman DM. ApoE4 markedly exacerbates tau-mediated neurodegeneration in a mouse model of tauopathy. Nature. 2017 Sep 28;549(7673):523–527. doi: 10.1038/nature24016. Epub 2017 Sep 20. PMID: 28959956; PMCID: PMC5641217.

32. Etkin A. A Reckoning and Research Agenda for Neuroimaging in Psychiatry. Am J Psychiatry. 2019 Jul 1;176(7):507–511. doi: 10.1176/appi.ajp.2019.19050521. PMID: 31256624.

33. Hao W, Friedman A. Mathematical model on Alzheimer’s disease. BMC Syst Biol. 2016 Nov 18;10(1):108. doi: 10.1186/s12918-016-0348-2. PMID: 27863488; PMCID: PMC5116206.

34. Puri IK, Li L. Mathematical modeling for the pathogenesis of Alzheimer’s disease. PLoS One. 2010 Dec 14;5(12):e15176. doi: 10.1371/journal.pone.0015176. PMID: 21179474; PMCID: PMC3001872.

35. Zhang Y, Chen H, Li R, Sterling K, Song W. Amyloid β-based therapy for Alzheimer’s disease: challenges, successes and future. Signal Transduct Target Ther. 2023 Jun 30;8(1):248. doi: 10.1038/s41392-023-01484-7. PMID: 37386015; PMCID: PMC10310781.

36. Arriagada PV, Growdon JH, Hedley-Whyte ET, Hyman BT. Neurofibrillary tangles but not senile plaques parallel duration and severity of Alzheimer’s disease. Neurology. 1992 Mar;42(3 Pt 1):631–9. doi: 10.1212/wnl.42.3.631. PMID: 1549228.

37. Arendt T, Stieler JT, Holzer M. Tau and tauopathies. Brain Res Bull. 2016 Sep;126(Pt 3):238–292. doi: 10.1016/j.brainresbull.2016.08.018. Epub 2016 Sep 9. PMID: 27615390.

38. Raj, A., Tora, V., Gao, X., Cho, H., Choi, J. Y., Ryu, Y. H., Lyoo, C. H., & Franchi, B. (2021). Combined Model of Aggregation and Network Diffusion Recapitulates Alzheimer’s Regional Tau-Positron Emission Tomography. Brain connectivity, 11(8), 624–638. 10.1089/brain.2020.0841

39. Cho H, Choi JY, Hwang MS, Kim YJ, Lee HM, Lee HS, Lee JH, Ryu YH, Lee MS, Lyoo CH. In vivo cortical spreading pattern of tau and amyloid in the Alzheimer disease spectrum. Ann Neurol. 2016 Aug;80(2):247–58. doi: 10.1002/ana.24711. Epub 2016 Jul 8. PMID: 27323247.

40. Gorgolewski K, Burns CD, Madison C, Clark D, Halchenko YO, Waskom ML, Ghosh SS. Nipype: a flexible, lightweight and extensible neuroimaging data processing framework in python. Front Neuroinform. 2011 Aug 22;5:13. doi: 10.3389/fninf.2011.00013. PMID: 21897815; PMCID: PMC3159964.

41. Jenkinson M, Bannister P, Brady M, Smith S. Improved optimization for the robust and accurate linear registration and motion correction of brain images. Neuroimage. 2002 Oct;17(2):825–41. doi: 10.1016/s1053-8119(02)91132-8. PMID: 12377157.

42. Smith JA, Knight RG. Memory processing in Alzheimer’s disease. Neuropsychologia. 2002;40(6):666–82. doi: 10.1016/s0028-3932(01)00137-3. PMID: 11792406.

43. McNab JA, Edlow BL, Witzel T, Huang SY, Bhat H, Heberlein K, Feiweier T, Liu K, Keil B, Cohen-Adad J, Tisdall MD, Folkerth RD, Kinney HC, Wald LL. The Human Connectome Project and beyond: initial applications of 300 mT/m gradients. Neuroimage. 2013 Oct 15;80:234–45. doi: 10.1016/j.neuroimage.2013.05.074. Epub 2013 May 24. PMID: 23711537; PMCID: PMC3812060.

44. Abdelnour F, Voss HU, Raj A. Network diffusion accurately models the relationship between structural and functional brain connectivity networks. Neuroimage. 2014 Apr 15;90:335–47. doi: 10.1016/j.neuroimage.2013.12.039. Epub 2013 Dec 30. PMID: 24384152; PMCID: PMC3951650.

45. Owen JP, Li YO, Ziv E, Strominger Z, Gold J, Bukhpun P, Wakahiro M, Friedman EJ, Sherr EH, Mukherjee P. The structural connectome of the human brain in agenesis of the corpus callosum. Neuroimage. 2013 Apr 15;70:340–55. doi: 10.1016/j.neuroimage.2012.12.031. Epub 2012 Dec 23. PMID: 23268782; PMCID: PMC4127170.

46. Verma P, Nagarajan S, Raj A. Spectral graph theory of brain oscillations--Revisited and improved. Neuroimage. 2022 Apr 1;249:118919. doi: 10.1016/j.neuroimage.2022.118919. Epub 2022 Jan 17. PMID: 35051584; PMCID: PMC9506601.

47. Zhu B, Liu Y, Hwang S, Archuleta K, Huang H, Campos A, Murad R, Piña-Crespo J, Xu H, Huang TY. Trem2 deletion enhances tau dispersion and pathology through microglia exosomes. Mol Neurodegener. 2022 Sep 2;17(1):58. doi: 10.1186/s13024-022-00562-8. PMID: 36056435; PMCID: PMC9438095.

48. Long H, Simmons A, Mayorga A, Burgess B, Nguyen T, Budda B, Rychkova A, Rhinn H, Tassi I, Ward M, Yeh F, Schwabe T, Paul R, Kenkare-Mitra S, Rosenthal A. Preclinical and first-in-human evaluation of AL002, a novel TREM2 agonistic antibody for Alzheimer’s disease. Alzheimers Res Ther. 2024 Oct 23;16(1):235. doi: 10.1186/s13195-024-01599-1. PMID: 39444037; PMCID: PMC11515656.

49. Lee SH, Bae EJ, Park SJ, Lee SJ. Microglia-driven inflammation induces progressive tauopathies and synucleinopathies. Exp Mol Med. 2025 May;57(5):1017–1031. doi: 10.1038/s12276-025-01450-z. Epub 2025 May 1. PMID: 40307569; PMCID: PMC12130470.

50. Oken RJ, McGeer PL. Schizophrenia, Alzheimer’s disease, and anti-inflammatory agents. Schizophr Bull. 1996;22(1):1–4. doi: 10.1093/schbul/22.1.1. PMID: 8685652.

51. Fan Z, Brooks DJ, Okello A, Edison P. An early and late peak in microglial activation in Alzheimer’s disease trajectory. Brain. 2017 Mar 1;140(3):792–803. doi: 10.1093/brain/aww349. PMID: 28122877; PMCID: PMC5837520.

52. Maphis N, Xu G, Kokiko-Cochran ON, Jiang S, Cardona A, Ransohoff RM, Lamb BT, Bhaskar K. Reactive microglia drive tau pathology and contribute to the spreading of pathological tau in the brain. Brain. 2015 Jun;138(Pt 6):1738–55. doi: 10.1093/brain/awv081. Epub 2015 Mar 31. PMID: 25833819; PMCID: PMC4542622.

53. Luo W, Liu W, Hu X, Hanna M, Caravaca A, Paul SM. Microglial internalization and degradation of pathological tau is enhanced by an anti-tau monoclonal antibody. Sci Rep. 2015 Jun 9;5:11161. doi: 10.1038/srep11161. PMID: 26057852; PMCID: PMC4460904.

54. Mezias C, LoCastro E, Xia C, Raj A. Connectivity, not region-intrinsic properties, predicts regional vulnerability to progressive tau pathology in mouse models of disease. Acta Neuropathol Commun. 2017 Aug 14;5(1):61. doi: 10.1186/s40478-017-0459-z. PMID: 28807028; PMCID: PMC5556602.

55. Bougacha S, Roquet D, Landeau B, Saul E, Naveau M, Sherif S, Bejanin A, Dhenain M, Raj A, Vivien D, Chetelat G. Contributions of connectional pathways to shaping Alzheimer’s disease pathologies. Brain Commun. 2025 Jan 6;7(1):fcae459. doi: 10.1093/braincomms/fcae459. PMID: 39763634; PMCID: PMC11702304.

56. Spires-Jones TL, Hyman BT. The intersection of amyloid beta and tau at synapses in Alzheimer’s disease. Neuron. 2014 May 21;82(4):756–71. doi: 10.1016/j.neuron.2014.05.004. PMID: 24853936; PMCID: PMC4135182.

57. Jackson RJ, Rudinskiy N, Herrmann AG, Croft S, Kim JM, Petrova V, Ramos-Rodriguez JJ, Pitstick R, Wegmann S, Garcia-Alloza M, Carlson GA, Hyman BT, Spires-Jones TL. Human tau increases amyloid β plaque size but not amyloid β-mediated synapse loss in a novel mouse model of Alzheimer’s disease. Eur J Neurosci. 2016 Dec;44(12):3056–3066. doi: 10.1111/ejn.13442. Epub 2016 Nov 12. PMID: 27748574; PMCID: PMC5215483.

58. Torok J, Mezias C, Raj A. Directionality bias underpins divergent spatiotemporal progression of Alzheimer-related tauopathy in mouse models. Alzheimers Dement. 2025 May;21(5):e70092. doi: 10.1002/alz.70092. PMID: 40396482; PMCID: PMC12093255.

59. Mancuso R, Van Den Daele J, Fattorelli N, Wolfs L, Balusu S, Burton O, Liston A, Sierksma A, Fourne Y, Poovathingal S, Arranz-Mendiguren A, Sala Frigerio C, Claes C, Serneels L, Theys T, Perry VH, Verfaillie C, Fiers M, De Strooper B. Stem-cell-derived human microglia transplanted in mouse brain to study human disease. Nat Neurosci. 2019 Dec;22(12):2111–2116. doi: 10.1038/s41593-019-0525-x. Epub 2019 Oct 28. PMID: 31659342; PMCID: PMC7616913.

60. Paolicelli RC, Sierra A, Stevens B, Tremblay ME, Aguzzi A, Ajami B, Amit I, Audinat E, Bechmann I, Bennett M, Bennett F, Bessis A, Biber K, Bilbo S, Blurton-Jones M, Boddeke E, Brites D, Brône B, Brown GC, Butovsky O, Carson MJ, Castellano B, Colonna M, Cowley SA, Cunningham C, Davalos D, De Jager PL, de Strooper B, Denes A, Eggen BJL, Eyo U, Galea E, Garel S, Ginhoux F, Glass CK, Gokce O, Gomez-Nicola D, González B, Gordon S, Graeber MB, Greenhalgh AD, Gressens P, Greter M, Gutmann DH, Haass C, Heneka MT, Heppner FL, Hong S, Hume DA, Jung S, Kettenmann H, Kipnis J, Koyama R, Lemke G, Lynch M, Majewska A, Malcangio M, Malm T, Mancuso R, Masuda T, Matteoli M, McColl BW, Miron VE, Molofsky AV, Monje M, Mracsko E, Nadjar A, Neher JJ, Neniskyte U, Neumann H, Noda M, Peng B, Peri F, Perry VH, Popovich PG, Pridans C, Priller J, Prinz M, Ragozzino D, Ransohoff RM, Salter MW, Schaefer A, Schafer DP, Schwartz M, Simons M, Smith CJ, Streit WJ, Tay TL, Tsai LH, Verkhratsky A, von Bernhardi R, Waxke H, Wittamer V, Wolf SA, Wu LJ, Wyss-Coray T. Microglia states and nomenclature: A field at its crossroads. Neuron. 2022 Nov 2;110(21):3458–3483. doi: 10.1016/j.neuron.2022.10.020. PMID: 36327895; PMCID: PMC9999291..ncbi.nlm.nih.gov/books/NBK27944/

